# Active site tyrosine residues in human NQO1 homodimer are critical for non-synchronous enzyme catalysis at the two active sites

**DOI:** 10.1101/2025.02.02.636097

**Authors:** Maribel Rivero, Juan Luis Pacheco-Garcia, Pavla Vankova, Dmitry S Loginov, Isabel Quereda-Moraleda, Jose Manuel Martin-Garcia, Petr Man, Milagros Medina, Angel Luis Pey

**Affiliations:** Department of Biochemistry and Molecular and Cellular Biology, Faculty of Sciences, University of Zaragoza, 50009 Zaragoza, Spain; Institute for Biocomputation and Physics of Complex Systems (BIFI), University of Zaragoza, 50018 Zaragoza, Spain; Department of Physical Chemistry, Faculty of Sciences, University of Granada, 18071, Granada, Spain; Institute of Biotechnology - BioCeV, Academy of Sciences of the Czech Republic, Vestec, Czech Republic; Institute of Microbiology - BioCeV, Academy of Sciences of the Czech Republic, Vestec, Czech Republic; Department of Crystallography and Structural Biology, Institute of Physical Chemistry Blas Cabrera, Spanish National Research Council (CSIC), Madrid 28006, Spain; Department of Physical Chemistry, Unit of Excellence in Chemistry applied to Biomedicine and Environment and Institute of Biotechnology, University of Granada, 18071, Granada, Spain

**Keywords:** Hydride transfer, Coenzyme specificity, Quinone oxidoreductase, Enzyme kinetics, Flavin

## Abstract

Human NQO1 is a flavoenzyme essential for the redox metabolization of many substances and associated with wide-impacting diseases such as cancer and Alzheimeŕs. Recent X-ray crystallographic studies have proposed that a few residues at the active site of NQO1 (including Tyr126 and Tyr128) may control enzyme catalysis and functional negative cooperativity. In this work, we use rapid mixing pre-steady state kinetics and hydrogen-deuterium exchange followed by mass spectrometry (HDX-MS) to evaluate experimentally the role of Tyr126 and Tyr128 in NQO1 functionality by generating mutants to Phe, Ala and Glu. Mutations to Phe caused mild effects, whereas those to Ala significantly decreased hydride transfer efficiency and those to Glu virtually abolished NQO1 activity. Interestingly, structural stability studies by HDX-MS showed significant perturbations particularly affecting the binding site of NADH/NAD^+^ in the less conservative mutations (particularly to Glu). Mutations of Tyr126 and Tyr128 seem to also modulate the non-synchronous catalysis in the two active sites (negative cooperativity) as well as the selectivity for NADH/NADPH as coenzymes. Our work experimentally demonstrates the critical role of Tyr126 and Tyr128 in the flavin reductive half-reaction of the catalytic cycle of NQO1 in the negative cooperativity, and also suggests that phosphorylation of these two Tyr residues might shut down NQO1 activity reversibly.

## 1. Introduction

NAD(P)H:quinone oxidoreductase 1 (NQO1; UniProt ID P15559) is a homodimeric flavoenzyme that plays a pivotal role in cellular defense mechanisms against oxidative stress and toxic quinones, contributing to control cellular redox homeostasis. It is a two-electron reductase that utilizes NADH or NADPH as hydride donors to catalyze the reduction of quinones to hydroquinones [1,2]. This reaction is essential for detoxification, as it prevents the formation of reactive oxygen species (ROS) through redox cycling. NQO1 is ubiquitously expressed in various tissues and is regulated by several pathways, including the antioxidant response element (ARE) mediated by the transcription factor NRF2 [1,2]. Its physiological functions extend beyond detoxification since it also stabilizes key proteins such as p53, p73 and HIF1α [1,3]. Additionally, NQO1 has garnered significant interest in cancer research due to its overexpression in certain tumors, making it a potential biomarker and therapeutic target [4].

Each monomer of NQO1 binds one flavin adenine dinucleotide (FAD) molecule as cofactor, thus the enzyme contains two active sites located near the FAD-binding region, where it has been shown to be able to accommodate coenzymes such as the hydride donors NADH or NADPH, and the hydride acceptor quinones [2,4]. Residues from both monomers contribute to make each of the two, in principle *identical*, active sites in NQO1. This configuration may facilitate two-electron reductions by hydride transfer (HT), thus reducing quinones to hydroquinones [5,6]. This mechanism also prevents redox cycling, minimizes oxidative stress and contrasts with one-electron reductions that promote the generation of ROS. Notably, pre-steady enzyme kinetic analysis of the human NQO1 (named along the manuscript as NQO1) revealed the existence of two different pathways for the FAD HT processes in both the flavin reductive and oxidative half-reactions, indicating that the reduction of the two FAD molecules within the NQO1 homodimer occurred at different rates thus supporting the existence of functional negative cooperativity [7–9]. Furthermore, room temperature crystallographic structures as well as molecular dynamics simulations evaluated for the interaction of NQO1 with NADH, clearly supported one of the two active sites of the homodimer preferentially holding the coenzyme in catalytically competent conformation over the other [6], further explaining cooperative effects observed for FAD, NADH and inhibitors binding [10–12].

Understanding the structure, mechanism, and regulation of NQO1 is vital for elucidating its roles in health and disease [4]. Structural studies pointed to a set of residues collectively orchestrating the NQO1 catalysis. Among them, Tyr126 and Tyr128 were particularly proposed to play critical roles in stabilizing the substrate and intermediates during the reduction process [5,6]. They were thought to participate in hydrogen bonding with the substrates (Figure 1), guiding their optimal allocation within the active site for catalysis. Understanding NQO1 catalytic mechanism is crucial to develop more efficient inhibitors for different approaches to treat cancer [13]. A recent study has also shown Tyr126 and Tyr128 may have been evolutionarily selected to provide NQO1 for its coenzyme specificity towards NADH and NADPH [14].

**Figure 1.**
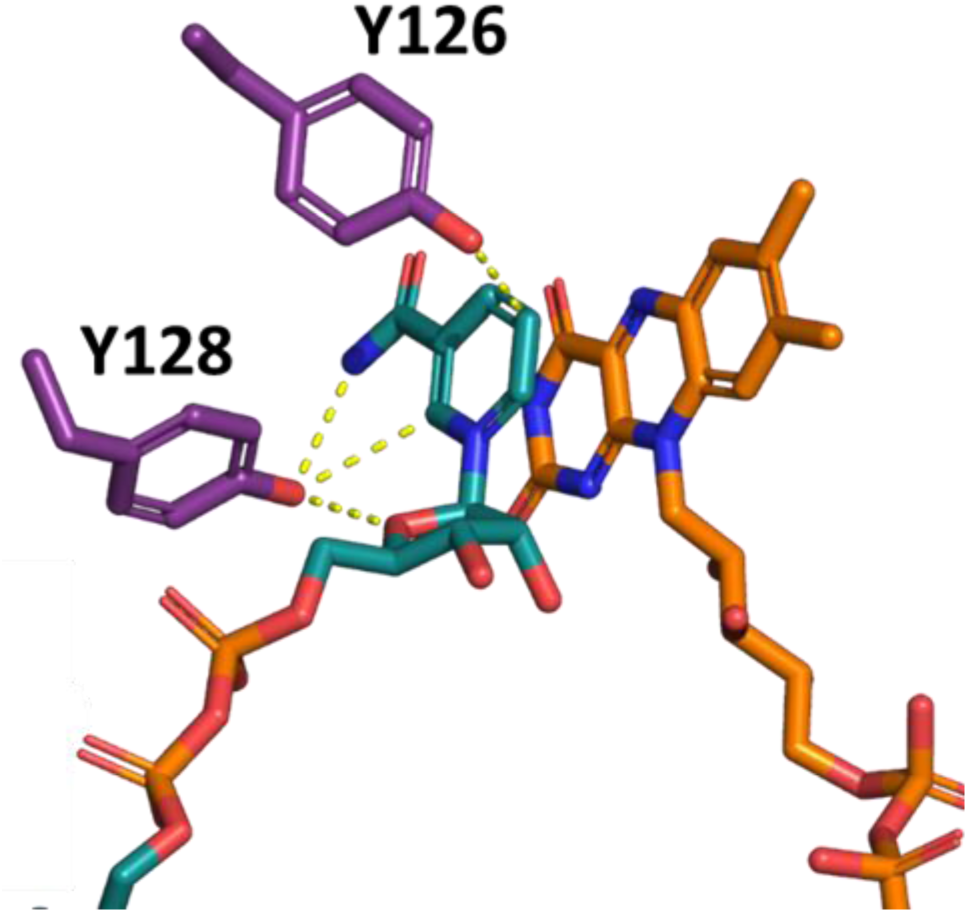
Structural arrangement of Tyr126 and Tyr128 with NAD^+^. Yellow dashed lines indicate hydrogen bonds between the two Tyr and the coenzyme. The structure used has generated using the PDB code 8RFM.

In this work, we experimentally investigate in-depth the role of Tyr126 and Tyr128 on NQO1 structure and function, by conservatively mutating them to Phe and non-conservatively to Ala. Mutations to Glu were also analyzed to probe the active site perturbations and to partially mimic the effect of phosphorylation in these two residues described by high throughput methods (as found in PhosphositePlus®, https://www.phosphosite.org/proteinAction.action?id=14721&showAllSites=true). We found that mutations to Ala noticeably affected NQO1 activity and cofactor specificity. The mutations to Glu virtually abolished NQO1 activity, thus suggesting that phosphorylation of these two tyrosine residues can be crucial to reversibly shut down NQO1 activity under physiological and pathological conditions.

## 2. Results and discussion

### 2.1. The rationale for generating mutants at Y126 and Y128

We have generated several mutants at positions 126 and 128 which are very close to the nicotinamide redox active moiety of NAD^+^/H (Figure 1). Two mutations were quite conservative (Y126F and Y128F), whereas four were much more disruptive, almost fully removing the bulky side-chain (Y126A and Y128A) or introducing a negative charge with a small change in side-chain size (Y126E and Y128E). It must be noted that mutants Y126E and Y128E might partially mimic phosphorylation events reported but not analysed in any detail (https://www.cellsignal.com/learn-and-support/phosphositeplus-ptm-database?srsltid=AfmBOoqkAo1yKVAIYlWjgFjdohbAmrtVX3u5sS2zF9ZtrJnAw-DOfTTJ).

All NQO1 variants were produced at good yields and isolated to a high purity (Figure S1). As purified, all of them showed the typical visible spectra for an almost saturated form of NQO1 with FAD, except for the variant Y126A (Figure S2). All NQO1 variants showed a wild-type (WT) like thermal stability, only with a small decrease in *T*_m_ for Y126A (Figure S3). Since FAD binding is linked to the thermal stability of NQO1 [12], the lower stability of Y126A might partially reflect a lower affinity for FAD in this variant. This notion is consistent with our visible absorption measurements (Figure S2)

### 2.2 Mutations of Y126 and Y128 affect the local stability of the holo-protein

HDX-MS has been instrumental in quantifying the effect of mutations and ligand binding on the structural stability and function of NQO1 [9,15–17]. The HDX behavior of NQO1 followed an EX2 mechanism in which the observed exchange rate constants reflect in part the local thermodynamic stability of the exchanging segments [15]. In most cases, the exchanging transients were well described by a single exponential function typically with a significant burst-phase (Figure S4-S8 and Table S1). This is an indication of significant conformational diversity in the native state ensemble of NQO1 with FAD bound (holo-NQO1, named as NQO1 along the manuscript), likely associated with the cooperative functional behavior towards substrates, cofactors, coenzymes and inhibitors [8,12,15,18].

We first used HDX-MS to quantify the effects of mutations at Y126 and Y128 on the local stability of NQO1 (in the holo form, *saturated* with FAD). The most remarkable effects were found in the Y126A and Y126E variants, while the behaviour of Y126F and Y128F was WT-like (Figure 2). In these two variants, regions 42-90, 105-112, 119-124, 129-141, 231-240 and 262-273 were thermodynamically destabilized according to an EX2 mechanism [15], particularly segments 54-74 and 131-140 (with Δ%D_av_ in the range of 20-40). Destabilization of the N-terminal segment 54-74 was confirmed by proteolysis experiments with thermolysin that cleaves between residues 71-72 [19], and showed a remarkable local thermodynamic destabilization by the mutations Y126A and Y126E of this region (in the range of 1.4-1.6 kcal·mol^-1^) whereas the mutant Y128A showed a milder destabilization (∼0.5 kcal·mol^-1^) (Figure 3 and S9, Table S2). The mutations Y126F, Y128E and Y128F had minimal effects on the local stability of NQO1 (Figure 3 and S9, Table S2).

**Figure 2.**
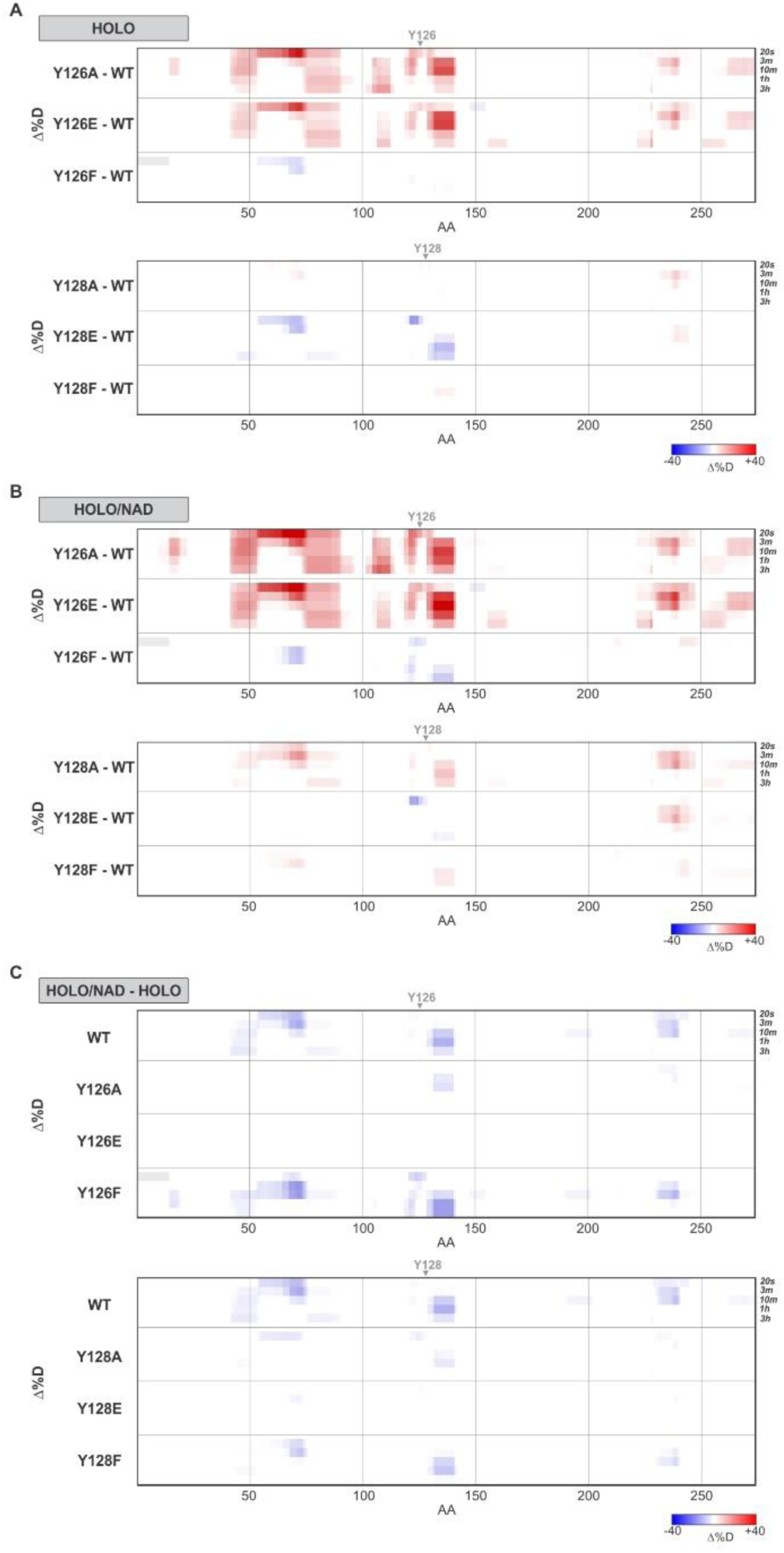
Differential heat maps for NQO1 WT and the Y126 and Y128 variants derived from HDX-MS. Destabilization (in shades of red) or protection (in shades of blue) (Δ%D) of the WT protein segments caused by mutations at Y126 (upper panel) and Y128 (lower panel) under NQO1 (A, HOLO, with an excess of FAD) or NQO1_NAD_ (B, with an excess of FAD and NAD^+^) conditions. The differences in deuterium incorporation for NQO1 variants in the presence or absence of NAD^+^ are shown in (C). Protein sequence is labelled on the x-axis. Exchange reaction times for corresponding rows are shown on the right. Color scale is shown under the plots.

**Figure 3.**
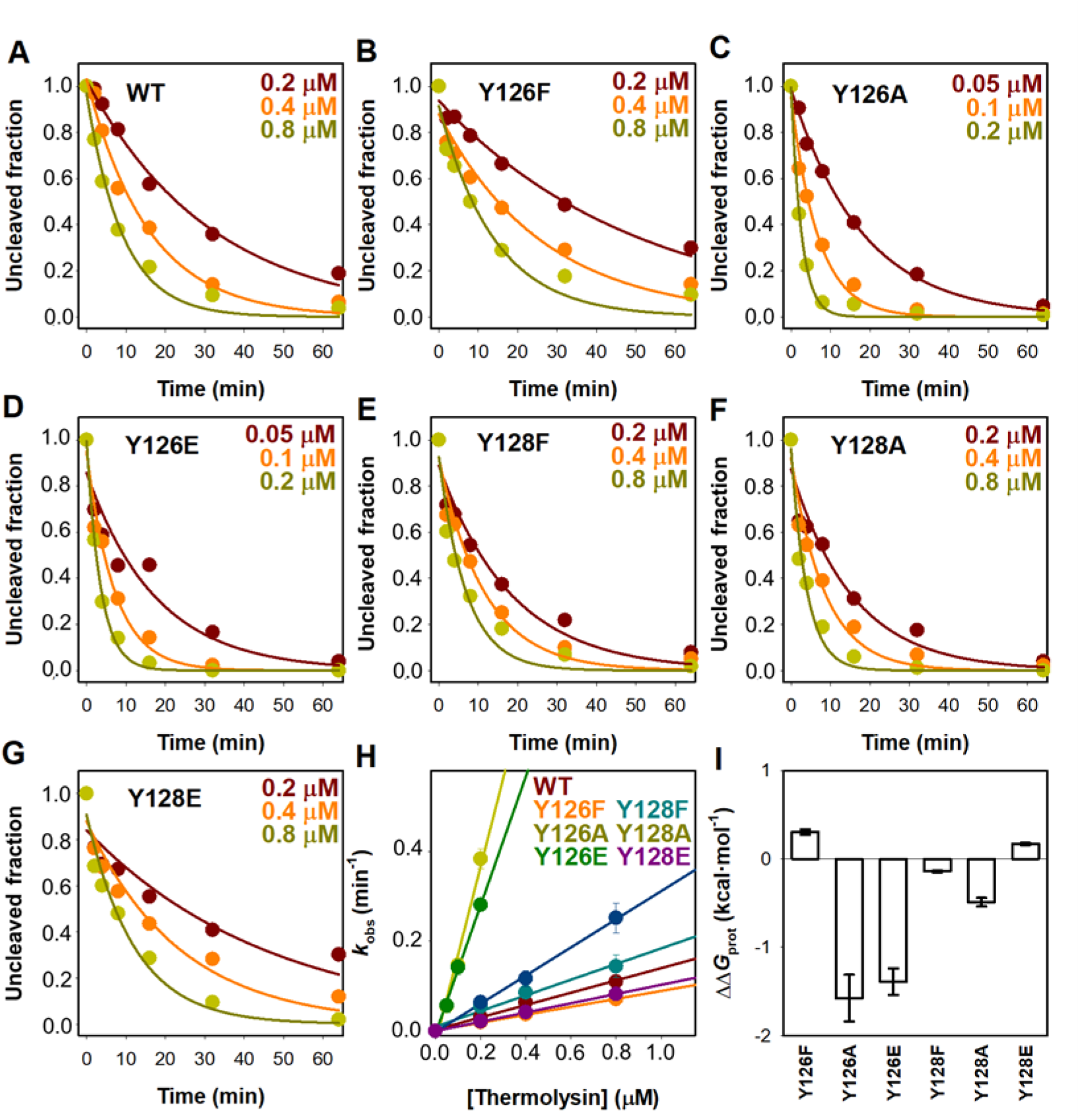
Proteolysis kinetics of NQO1 variants in the presence of thermolysin. Panels A-G show densitometric analysis of SDS-PAGE gels for the different NQO1 variants in the presence of different thermolysin concentrations (see color code in each panel; results were from a single experiment). Lines are best-fits to a single exponential function that yield the first-order rate constants (*k*_obs_). Panel H shows the dependence of *k*_obs_ on thermolysin concentration (the color code indicates the NQO1 variant analyzed and errors are those from fittings shown in panels A-G). Lines are best-fits to a linear function whose slopes provide the second-order rate constants *k*_prot_ (see table S2). Panel I shows the effect of mutations on the local thermodynamic stability in the vicinity of the primary cleavage site (S71-V72; [19]) from the values of *k*_prot_ as follows: ΔΔ*G*_prot_= - R·T·ln (*k*_prot(mutant)_ /(*k*_prot(WT)_).

### 2.3 NAD^+^ binding stabilizes WT NQO1 but not Y126E/A and Y128E/A mutants through long range effects

The effects of binding of NAD^+^ to NQO1 were also analyzed by HDX-MS (Figure 2 and 4, Figure S4-S9 and Table S1). In the presence of a large excess of NAD^+^, the WT protein showed stabilization (moderate, between 10-20 Δ%D_av_) of three segments, 65-75, 132-140 and 232-239. Using a cut-off of 5-10 Δ%D_av_, segments 42-64, 76-89, 121-124,

**Figure 4.**
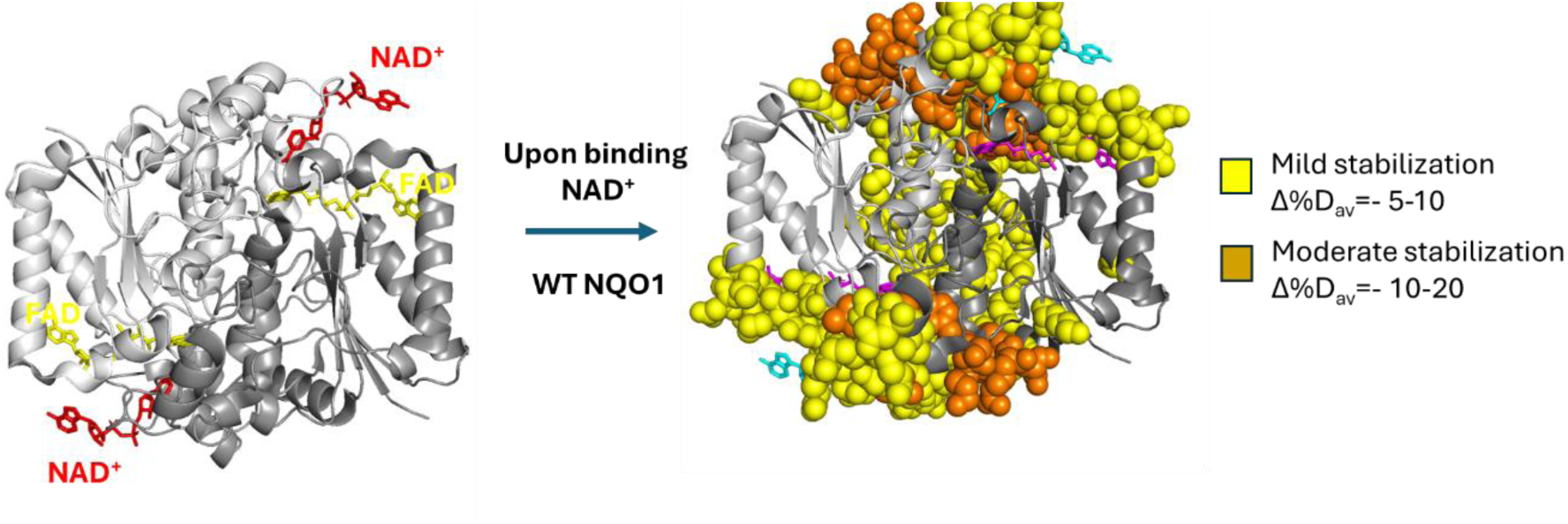
Binding of NAD^+^ globally stabilizes WT NQO1. Data were calculated from HDX-MS results. Calculations of Δ%D_av_ were done as described in [15,17]. The structural display is based on a symmetrical model generated from the structure of NQO1 with NAD^+^ bound [6].

129-131, 141, 191-201, 229-231, 240-244 and 269-270 were mildly stabilized (Figure 2 and 4). Noteworthy, these stabilizations do not only affect locally the binding site and long-range propagated (Figure 4). A similar effect was observed in the mutants Y126F and Y128F, but little or no stabilization was observed in the Y126E/A and Y128E/A, supporting altered NAD^+^ binding (possibly due to very low affinity) (Figure 2C). Consequently, the mutants Y126E/A and Y128E/A are *overdestabilized* vs. the WT protein at these segments in the presence of NAD^+^ (Figure 5 and S9). That also indicates that destabilization caused by the mutations Y126E/A and Y128E/A propagates far from the mutated sites (Figure 2A and 4).

**Figure 5.**
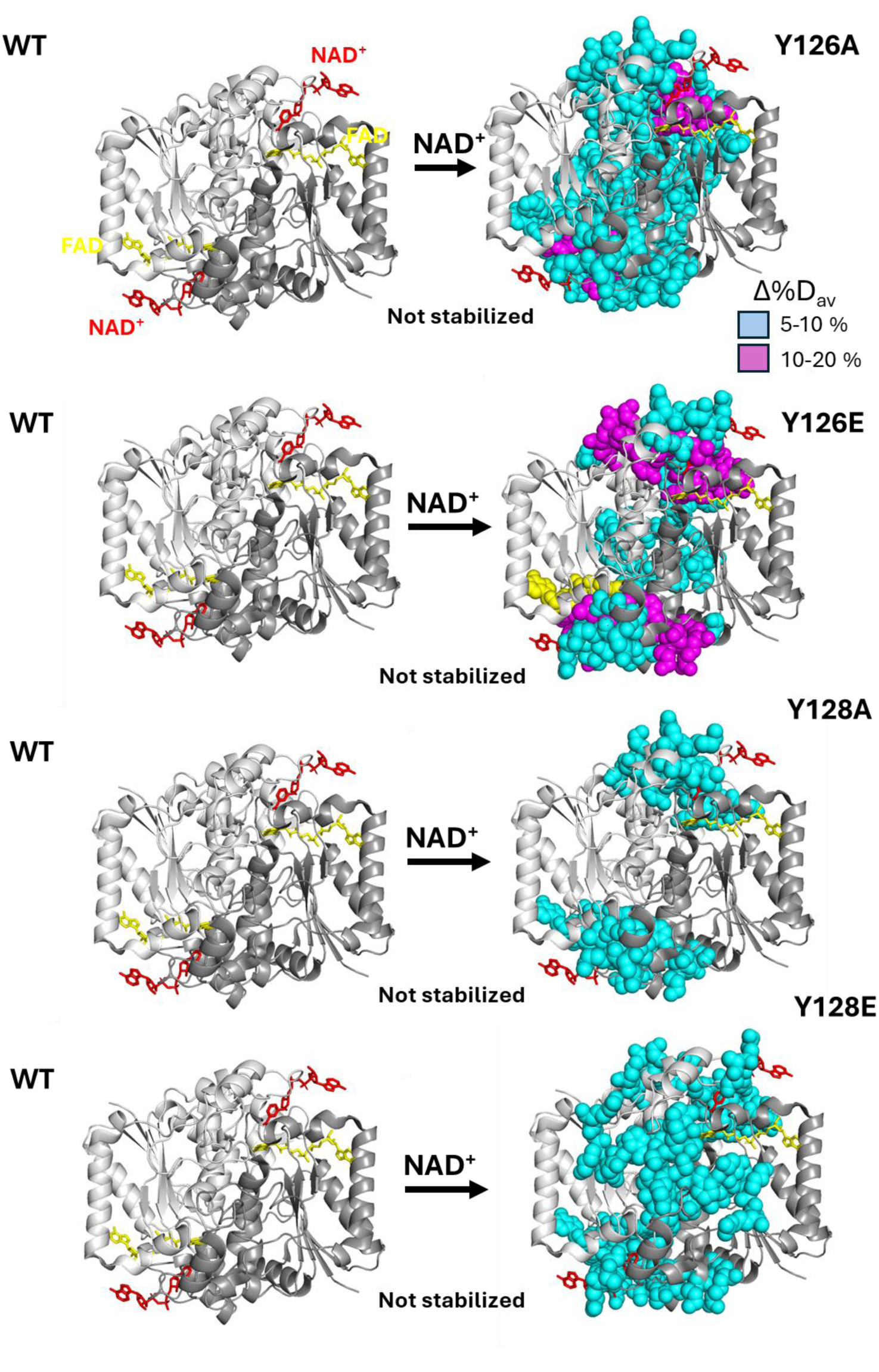
Mutations at Tyr126 and Tyr128 disrupt binding of NAD^+^. Calculations of Δ%D_av_ were done as described in [15,17] as the difference between a given mutant and the WT. The structural display is based on a symmetrical model generated from the structure of NQO1 with NAD^+^ bound [6].

### 2.4 The role of Tyr126 and Tyr128 on the NADH reductive half-reaction of NQO1

Pre-steady enzyme kinetic analysis of WT NQO1 has revealed the existence of two different pathways for the FAD reductive half-reaction by the NADH coenzyme, termed *fast* and *slow*, indicating that the reduction of the two FAD molecules within the NQO1 homodimer occurred at different rates [8]. Moreover, only the *fast* process was compatible with the catalytic constant, making also the flavin reductive half-reaction the limiting step in overall catalysis. When evaluating the NADH flavin reductive-half reaction for variants at positions Y126 and Y128 of NQO1, mutants specifically perturbed some features compared to the WT enzyme (Table 1 and Figure S11). Mutations affected times required for achieving reduction of each of the two FAD cofactors within the homodimer and/or ratio of oxidized/reduced FAD cofactors upon achieving the reaction equilibrium (Figure S11). Nonetheless, spectral deconvolutions of multi-wavelength time-resolved data indicated that for all variants reduction of the FAD cofactors occurs through two steps as described for the WT enzyme [8] (Figure S12). Noticeably, the deleterious effects caused by some of the mutations in the *k*_obs,fast_ values (please, see Table 1) made also possible to experimentally follow the FAD reductive-half reduction using as hydride donor NADPH at different coenzyme concentrations, a process that was too fast to be evaluated for the WT as well as for some of the variants (Figures S13 and S14).

**Table 1.** Kinetic parameters for the reductive half-reaction of the NQO1 variants at Y126 and Y128 with NADH and NADPH. Primary data and fittings are shown in Figure 4.

| *NQO1 REDUCTION BY NADH* |  |  |  |  |  |  |
| --- | --- | --- | --- | --- | --- | --- |
|  | *Fast* |  |  | *Slow* |  |  |
| | $k_{HT}$<br>(s<sup>-1</sup>) | $K_d^{NADH}$<br>(μM) | $k_{HT}/K_d^{NADH}$<br>(s<sup>-1</sup>·μM<sup>-1</sup>) | $k_{HT}$<br>(s<sup>-1</sup>) | $K_d^{NADH}$<br>(μM) | $k_{HT}/K_d^{NADH}$<br>(s<sup>-1</sup>·μM<sup>-1</sup>) |
| WT<sup>a</sup> | 281±12 | 15.2±1.9 | 18.5±2.4 | 14.3±1.5 | 8.2±3.3 | 1.7±0.7 |
| Y126A | 50±2 | 32±2 | 1.7±0.3 | 24±2 | 45±10 | 5.3±1.5·10<sup>-1</sup> |
| Y128A | 53±1 | 19±1 | 2.8±0.2 | 9.3±0.8 | 10.4±4.4 | 8.9±4.5·10<sup>-1</sup> |
| Y126E | 1.8±0.4·10<sup>-1</sup> <sup>b</sup> | N.a. <sup>b</sup> | N.a. <sup>b</sup> | 2.8±1.4·10<sup>-2</sup> <sup>b</sup> | N.a. <sup>b</sup> | N.a. <sup>b</sup> |
| Y128E | 9.7±0.8·10<sup>-2</sup> <sup>c</sup> | N.a. <sup>c</sup> | N.a. <sup>c</sup> | 3.9±0.2·10<sup>-2</sup> <sup>c</sup> | N.a. <sup>c</sup> | N.a. <sup>c</sup> |
| Y126F | 290±12 | 28±3 | 10.3±1.5 | 16.1±1.5 | 33±5 | 4.9±1.8·10<sup>-1</sup> |
| Y128F | 245±20 | 46±7 | 5.3±1.2 | 6.7±0.7 | 9.0±2 | 7.4±2.2·10<sup>-1</sup> |
| *NQO1 REDUCTION BY NADPH* |  |  |  |  |  |  |
|  | *Fast* |  |  | *Slow* |  |  |
| | $k_{HT}$<br>(s<sup>-1</sup>) | $K_d^{NADPH}$<br>(μM) | $k_{HT}/K_d^{NADPH}$<br>(s<sup>-1</sup>·μM<sup>-1</sup>) | $k_{HT}$<br>(s<sup>-1</sup>) | $K_d^{NADPH}$<br>(μM) | $k_{HT}/K_d^{NADPH}$<br>(s<sup>-1</sup>·μM<sup>-1</sup>) |
| WT<sup>a</sup> | 261±13 <sup>d</sup> |  |  | 7.8±0.3 <sup>b</sup> |  |  |
| Y126A | 40±1 | 6.3±1.4 |  | 18.4±0.9 | 17.2±2.0 |  |
| Y128A | 175±5 | 35±3 |  | 21.2±1.8 | 11.0±2.7 |  |
| Y126E | 3.2±0.1·10<sup>-3</sup> <sup>c</sup> | N.a. <sup>c</sup> | N.a. <sup>c</sup> | 9.2±2.0·10<sup>-4</sup> <sup>c</sup> | N.a. <sup>c</sup> | N.a. <sup>c</sup> |
| Y128E | 2.5±0.2·10<sup>-1</sup> <sup>c</sup> | N.a. <sup>c</sup> | N.a. <sup>c</sup> | 8±1 ·10<sup>-2</sup> <sup>c</sup> | N.a. <sup>c</sup> | N.a. <sup>c</sup> |
| Y126F | 78±6 <sup>d</sup> |  |  | 3.0±0.5 <sup>d</sup> |  |  |
| Y128F | 169±5 <sup>d</sup> |  |  | 12.5±0.9 <sup>d</sup> |  |  |

<sup>a</sup> Data from [8,20].
<sup>b</sup> $k_{obs}$ is independent on [NAD(P)H], therefore data do not allow to determine $K_d^{NAD(P)H}$ (N.a., not applicable) and the given value corresponds to a limiting $k_{HT}$ calculated as the average of $k_{obs}$ at different [NAD(P)H].
<sup>c</sup> $k_{obs}$ is linearly dependent on [NAD(P)H], therefore data do not allow to determine neither $K_d^{NAD(P)H}$ nor $k_{HT}$ (N.a., not applicable) and the given value corresponds to the second-order rate constant of the process calculated as the slope of the $k_{obs}$ vs. [NAD(P)H] plot (s<sup>-1</sup>·μM<sup>-1</sup>).
<sup>d</sup> Reaction is too fast and $k_{obs}$ values (particularly for the fast process, since it occurs in part within the instrumental dead time) cannot be accurately determined when increasing the NADPH:NQO1 ratio. The presented values correspond to $k_{obs}$ values determined at stoichiometric concentrations.

The variants Y126A or Y128A were able to become nearly fully reduced by both NADH and NADPH, particularly Y126A (Figure S11A-B and Figure S13A-B). Moreover, deconvolution of spectral evolutions showed that in both variants (and despite Y126A is not fully uploaded with FAD) the reduction process behaved similarly to the WT enzyme, resembling two processes, a *fast* and a *slow* [8,20], with the former showing just a slightly larger amplitude (representative analysis are shown in Figures S12A-B and S14A-B). Nonetheless, mutations had a particular negative impact on *k*_obs,fast_ (the observed rate constant for the *fast* process) values along the evaluated NADH and NADPH concentration ranges, while *k*_obs,slow_ ones remained in general closer to those for the WT (Figure 6 and Table 1). As for the WT enzyme, Y126A and Y128A *k*_obs,fast_ values showed a hyperbolic dependence on both NADH and NADPH coenzyme concentrations (Figure 6A and 6C) that allowed estimation of the dissociation constant (*K*_d,fastNAD(P)H_) for the NAD(P)H:NQO1 reactive complexes as well as a limiting value for the HT rate constant (*k*_HT,fastNAD(P)H_). Y126A and Y128A respectively increase *K*_d,fastNADH_ compared to the WT enzyme by 2.1 and 1.3 fold, while both mutants decreased *k*_HT,fastNADH_ by ∼ 5.5 fold (Table 1). These analyses revealed that these mutants have respectively only 10% and 15% of the catalytic efficiency exhibited by the WT in the *fast* process. These mutants showed milder effects on the *slow* process, respectively decreasing the efficiency to 30% and 55 % of that of WT (Table 1), indicating changes in synchronization at the two active sites (negative cooperativity). Regarding NADPH as donor, the reaction for the WT enzyme was too fast so a full comparison cannot be done. However, *K*d,fast^NADPH^ for Y126A and Y128A were respectively 5-fold lower and 1.8-fold higher regarding *K*d,fast^NADH^. On their side, while *k*HT,fast^NADPH^ for Y126A was nearly the same as *k*HT,fast^NADH^, in the case of Y128A this value increased by 3.3-fold. Overall, these data result in the *fast* process being more efficient (by 3.7 and 1.7 fold) for the variants Y126A and Y128A, with NADPH than with NADH. Impact of these mutations on the *slow* process, despite with apparent minor magnitude changes in *K*_d,fast_ and *k*HT,fast^NADPH^, also made this process being in both variants over 2-fold faster with NADPH than with NADH. Overall, these results support that these two highly non-conservative mutations retain the specificity towards NADPH, even though they affect the functional cooperativity at the two active sites possibly through changes in the native state ensemble (i.e. changes in *more competent* vs. *less competent* catalytic state ratios) [6,8].

**Figure 6.**
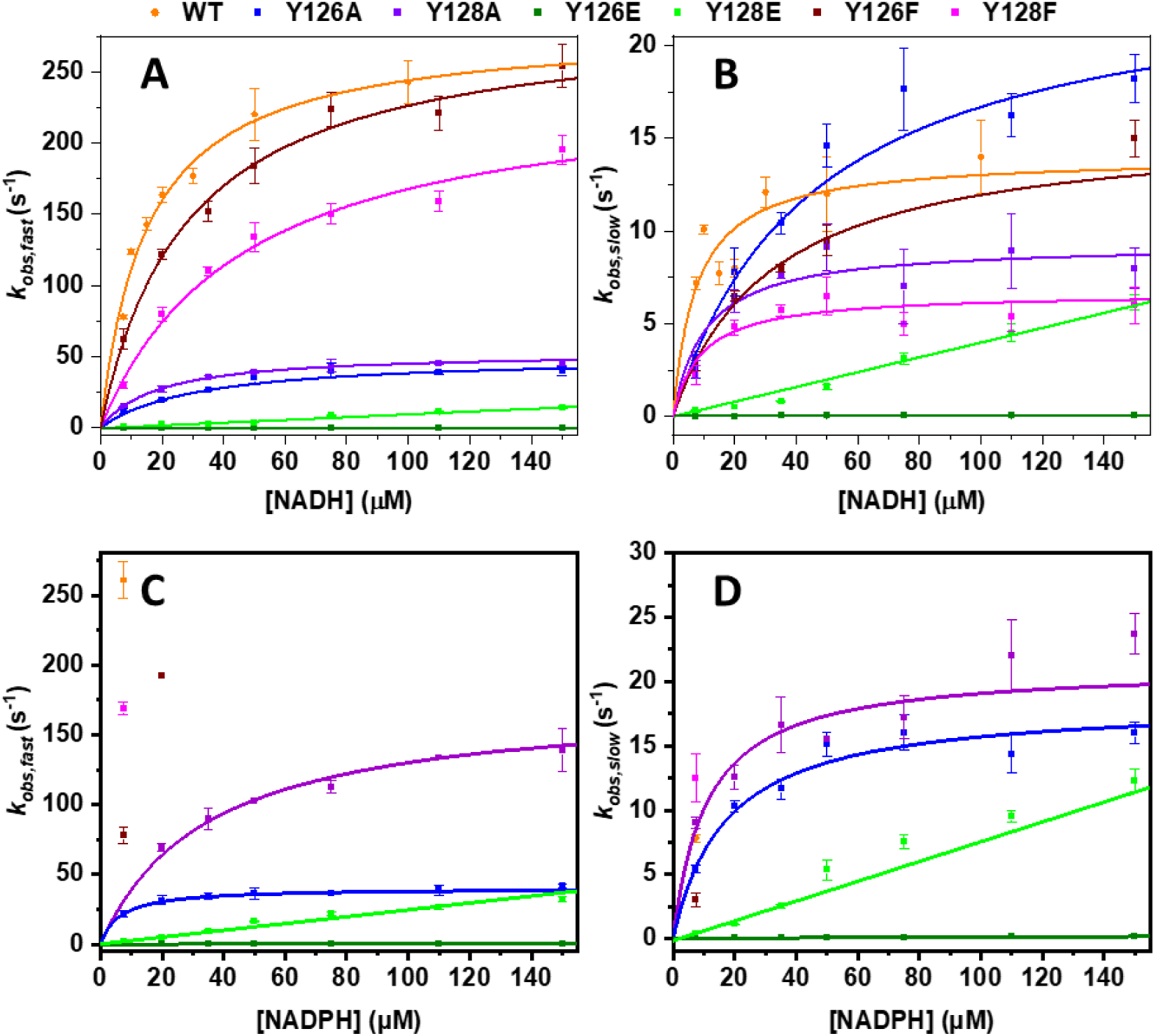
Impact of mutations at Y126 and Y128 of NQO1 on the pre-steady state kinetic parameters for the NAD(P)H dependent reductive half-reaction. Dependence of the observed rate constants for the (A) *fast* and (B) *slow* processes on NADH concentration, and for the (C) *fast* and (D) *slow* processes on NADPH concentration. Experimental data for each mutant are coloured by symbols as indicated in the figure code. Lines are best fits to a hyperbolic function or to a straight line. Errors are the s.d. from at least three replicates.

The Y126E and Y128E variants in NQO1 had a strong deleterious impact on the NAD(P)H dependent flavin reductive half reaction, at both the *fast* and *slow* processes (Table 1, Figures S11C-D and S12C-D, Figures S13C-D and S14C-D). Despite nearly full reduction of the FAD was achieved for both variants with both coenzymes, the corresponding values for *k*_obs,fast_ and *k*_obs,slow_ were very small as compared with those for the WT under equivalent conditions (Figure 6, Table 1). Noticeably, in this case the amplitude for the *fast* process resulted smaller than that of the *slow* process. These results suggest that the *slow* pathway is outweighing the *fast* one, and evidently affects the functional cooperativity at the two active sites of the enzyme. Moreover, while for both *fast* and *slow* processes *k*_obs_ ^NAD(P)H^ values of Y128E showed a very slight linear dependence on the NAD(P)H concentration (allowing to estimate a very slow second-order rate constant), the corresponding values for Y126E showed a lineal dependence on NADPH concentration but resulted in an essentially independence on the NADH concentration (Figure 6, Table 1). These results further supported a remarkable effect on the coenzyme performance when Y126 or Y128 are largely perturbed. Noticeably, *k*_obs_ ^NADPH^ values for both mutants were slightly larger than *k*_obs_ ^NADH^ ones in the concentration range evaluated. Altogether, these observations suggest that the introduced Glu prevents NQO1 from achieving the competent binding state for HT from the nicotinamide of the coenzyme to the isoalloxazine of FAD, thus limiting the overall kinetics [20]. These data also indicate that the deleterious impact in HT is larger when substituting Y126 as compared with Y128.

Remarkably, the conservative variants Y126F and Y128F produced NQO1 enzymes for which reduction of their FAD cofactor reached up to around 65-70% within the NAD(P)H concentration ranges evaluated (Table 1, Figures S11E-F and S13E-F). Moreover, deconvolution of spectral evolutions showed how, in both variants, the FAD reduction best fitted to a two steps process, *fast* and *slow*, with the first showing just a slightly larger amplitude (representative analysis are shown in Figures S11E-F and S13E- F). It is also worth to point out that in both mutants a final slowdecrease in the amplitude of the flavin band I is observed, independently of the coenzyme form. This process occurred after separated from the *fast* and *slow* steps with a lag phase, and shows very low *k*_obs_ values which are nearly independent on the NAD(P)H concentration. As for the WT, *k*obs,fast^NADH^ and *k* ^NADH^ showed hyperbolic dependences on the NADH concentration (Figure 6A-B), while when using NADPH as hydride donor the large *k*obs,fast^NADPH^ values prevented to evaluate the NADPH concentration profile for the reductive half reaction (Table 1). They showed minor effects in *k*HT,fast^NADH^ values, whereas for both variants the *K*d,fast^NADH^ mildly increased by ∼1.8 and ∼3 fold for Y126F and Y128F respectively, resulting in these mutants having 55% and 30% of the catalytic efficiency exhibited by the WT in the *fast* process. In the case of the *slow* process, these mutants respectively decreased the efficiency to 30% and 43 % of that of WT.

### 2.5. Mutations at Y126 affect the selectivity for NADH/NADPH as coenzymes and functional negative cooperativity

Despite the activity data for the set of NQO1 variants investigated is very complex to be analyzed in a simple manner (e.g. some reactions are too fast, some concentration dependencies are hyperbolic or linear, and so forth) (section 2.4), we aimed to analyze the coenzyme preference for the mutants at Tyr126 and Tyr128. To this end, we simply compared the observed rate constants for HT transfer for the *fast* and *slow* paths (Table 1). These analyses supported that the Y126E mutant clearly preferred NADH over NADPH as coenzyme (with a change of ∼30-50 fold in the rate constant for the *fast* and *slow* pathways) (Figure 7). We also observed a similar but milder effect for Y126F variant (∼4-5-fold increase for the *fast* and *slow* pathways). This contrasts with the clear preference for NADPH in the case of WT NQO1. These results clearly support the role of Tyr126 in coenzyme differential recognition [6,14].

**Figure 7.**
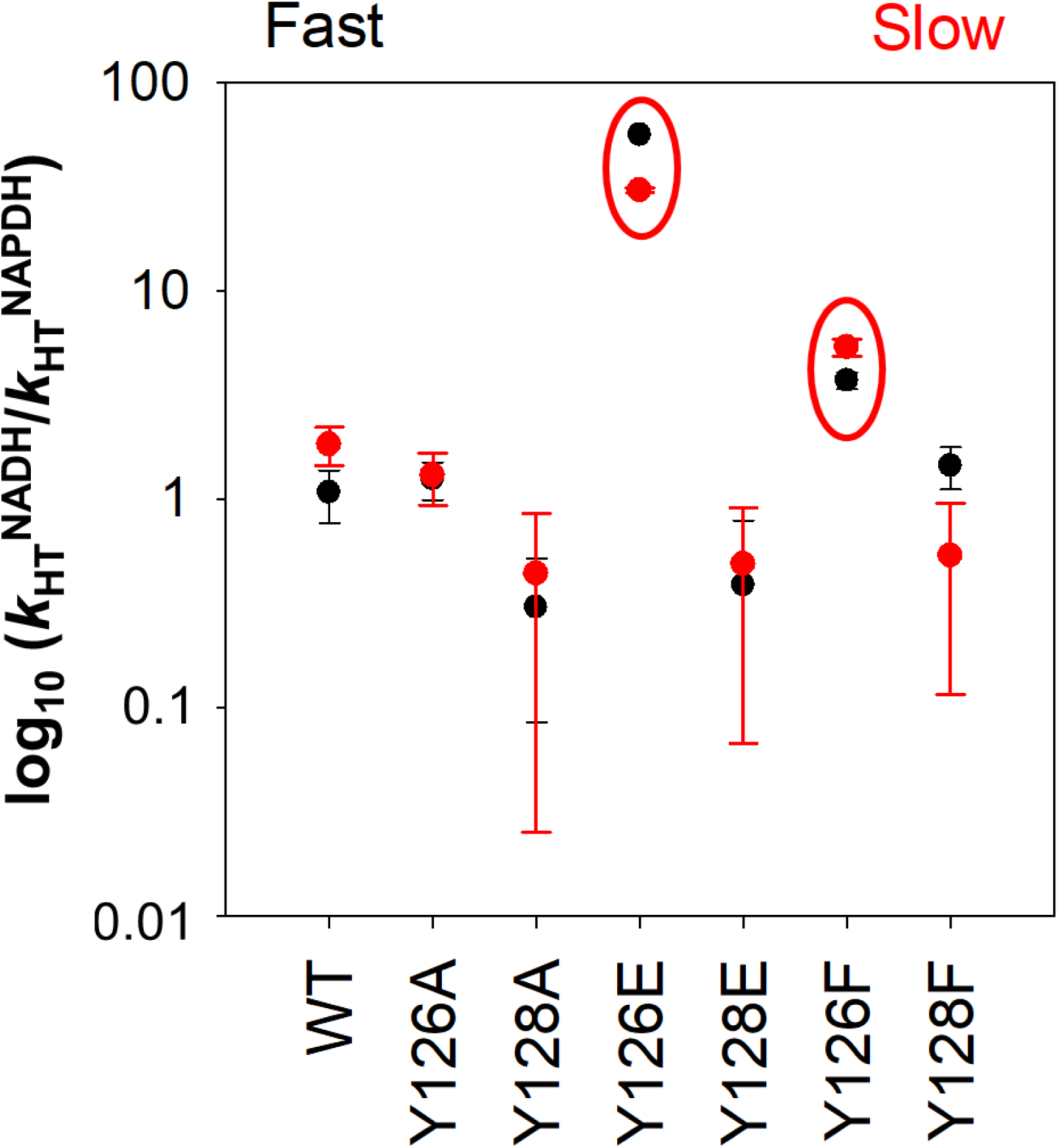
Mutations at Y126 change the selectivity between NADH and NADPH as coenzymes. The ratio of *k*_HT_ using NADH and NADPH were calculated from Table 1. Errors were linearly propagated from fitting errors. Red ellipsoids indicate those variants with a much larger preference for NADH.

The asynchronous active sites is likely the origin of NQO1 functional cooperativity [8,10]. A simple manner to evaluate this functional cooperativity is to compare the ratios of the *k*_cat_ for the fast and slow processes for the mutants and divide them by the value for the WT protein, that we named *cooperativity index* (data in Table 1). Using this index, four mutants affected the functional cooperativity. Two of them clearly reduced the functional cooperativity (Y126A, by 10- and 15-fold, using NADH and NADPH, respectively; Y128E, by 8- and 11-fold, using NADH and NADPH, respectively). A similar but milder effect was observed for Y128A (by 4-fold, using both NADH and NADPH). The mutation Y126E had different behaviors using NADH (cooperativity increased by 3-fold using NADH but decreased by 10-fold using NADPH).

### 2.6 Mutations at Y126 and Y128 hardly affect the NQO1 FAD reduction potential

NQO1 underwent reduction of its FAD by the non-physiological electron donor xanthine/xanthine oxidase (XO) system, with the indigo disulphonic acid two-electron exchanger being the dye that better equilibrated with FAD reduction and monitored by absorption spectroscopy (Figure S15A). The dependence of the logarithmic oxidized/reduced concentrations of the enzyme versus those of the dye showed a slope close to 1. This agreed with a two-electron exchange, and allowed to estimate a mid-point reduction potential, *E*_humanNQO1ox/hq_, of -125 mV (Figure S15B), being this value less negative than that of free FAD (*E*_FADox/hq_=-219 mV, [21], as well as than that reported for the NQO1 rat enzyme (*E*_ratNQO1ox/hq_ =-159 ± 3 mV) [22]. A similar procedure was also used to estimate *E*_NQO1ox/hq_ for the different variants at positions Y126 and Y128 (Figure S15B and Table S3). Mutations had minor impacts on *E*_humanNQO1ox/hq,_ with values being in general between 4 and 15 mV more negative than for the WT protein, Y126E NQO1 being the mutant exhibiting the more negative midpoint potential. Noticeably, during determination of these midpoint potentials some features appeared in the flavin band II that might be compatible with the transient stabilization of a flavin anionic semiquinone. To confirm this, photoreduction and reoxidation experiments were carried out (Figures S15C-D) and the appearance of anionic FAD semiquinone was observed upon reduction by photoirradiation. This state was not observed during the protein reoxidation by molecular oxygen, and it is not expected during the HT processes involving the action of this enzyme *in vivo*.

## 3. Conclusions

In this work, we have experimentally confirmed recent predictions on the key roles of Tyr126 and Tyr128 on enzyme catalysis, synchronization of active sites and preference for NADPH versus NADH as coenzymes in the human NQO1 homodimeric flavoenzyme [6,8,10,14,18]. We systematically described how large perturbations at these residues lead to dramatic drops in catalytic efficiency, changes in the catalytic mechanism and coenzyme selectivity through local structural destabilization that propagated in the long-range. Thus, our work provides at least three essential conclusions. First, the hydrogen bonding network between hydroxyls of Tyr126 and Tyr128 with NADH are not crucial for proper NQO1 catalysis (mutations to Phe). Second, development of inhibitors or activators of NQO1 for therapeutic purposes [4,13] must take into strong account the key roles in the NQO1 structure and dynamics of Tyr126 and Tyr128, NQO1 catalytic cycle and non-synchronization of redox processes at the two active sites. Third, the fact that these two Tyr residues undergo phosphorylation events (https://www.cellsignal.com/learn-and-support/phosphositeplus-ptm-database?srsltid=AfmBOooDTuV7LiI-MUWwLh_MKedODzpc9CV5HQ3FK0mR4OTmnBvh_gLo) [23], introducing negative charges (as we mimic in the Y126E and Y128E variants), may reversibly inactivate NQO1 function, with physiological and pathological implications. This may add another layer of complexity to understand the relationships between genotypic diversity of NQO1 and the propensity to develop disease [9,16,20,24–27]. In addition, our study shows that binding of NAD^+^ (and possibly NADH) causes mild stabilization propagating far beyond the binding site, whereas mutations Y126E/A and Y128E/A caused a moderate destabilization that propagates far from the mutated sites. This study thus confirm the large plasticity and response to perturbations (mutations, ligand binding, phosphorylation) observed in previous studies [6,7,15–17,20].

## 4. Materials and methods

### 4.1 Protein expression and purification

Mutations were introduced by site-directed mutagenesis in the WT NQO1 cDNA cloned into the pET-15b vector (pET-15b-NQO1) by GenScript (Leiden, The Netherlands). Codons were optimized for expression in *Escherichia coli* and mutagenesis was confirmed by sequencing the entire cDNA. The plasmids were transformed in *E. coli* BL21(DE3) cells (Agilent Technologies) for protein expression. These constructs contain a hexa-his N-terminal tag for purification. Expression and purification of NQO1 variants were carried out as described [8,15] using immobilized nickel affinity chromatography columns (Cytiva) and size-exclusion chromatography (Cytiva). Isolated dimeric fractions of NQO1 variants were buffer exchanged to HEPES-KOH buffer 50 mM pH 7.4 using PD-10 columns (Cytiva). The UV–visible spectra of purified NQO1 proteins were registered in a HP8453 UV–Visible spectrophotometer (Agilent Technologies,) and used to quantify the content of FAD as described [8] using protein variants from two different purifications.

### 4.2 Thermal denaturation

Thermal denaturation of NQO1 variants (2 μM of purified protein with added 20 μM FAD, from Merck) was monitored by following changes in tryptophan emission fluorescence in HEPES-KOH 50 mM at pH 7.4 as described [28]. *T*_m_ values were reported as mean ± s.d. of three independent measurements.

### 4.3 Proteolysis

Kinetic proteolysis studies by thermolysin were performed in 50 mM HEPES-KOH, 10 mM CaCl_2_ at pH 7.4 and 25 °C, as recently described [19,28]. 10-20 μM NQO1 variants as purified were incubated with an excess of 100 μM FAD, and experiments were carried out using a (final) thermolysin concentration of 0.05–0.8 μM. Proteolysis was triggered by addition of a concentrated protease stock (x10), samples were withdrawn at different time points and stopped with 25 mM EDTA (Ethylenediaminetetraacetic acid, Merck) at pH 8.0. Samples were denatured with Laemmlís buffer and analyzed by polyacrylamide gel electrophoresis in the presence of SDS (SDS-PAGE) containing 12% acrylamide. Gels were scanned and analyzed using the imageJ software (https://imagej.nih.gov/). Time-dependent decay of uncleaved NQO1 was used to determine first-order rate constants (*k*_obs_) from fittings carried out using an exponential function. Second-order rate constants (*k*_prot_) were obtained by dividing first-order rate constants by the protease concentration used (Table S2). Mutational effects on the local thermodynamic stability of NQO1 variants (ΔΔ*G*_prot_; primary cleavage site at S71-V72) were determined as previously described [9,29].

### 4.4 Hydrogen-deuterium exchange monitored by mass spectrometry (HDX-MS)

Hydrogen-deuterium exchange kinetics were monitored for all NQO1 variants under the FAD (NQO1_HOLO_) and FAD-NAD^+^ (NQO1_NAD_) conditions. Proteins were pre-incubated for 5 min at 21 °C with 10-molar excess of FAD prior to the exchange. In the case of NQO1_NAD_ conditions, another pre-incubation with 500-molar excess of NAD was done. HDX was carried out at 21 °C using a 10-fold dilution to D_2_O-based buffer (50 mM HEPES-KOH, pD 7.4/pH_read_ 7.0) enriched with 5 mM NAD^+^ for NQO1_NAD_ conditions. Protein concentration during exchange was 1 μM. Exchange times were 20, 180, 600, 3600 and 10800 s, and the time points 20, 180 and 3600 s were triplicated yielding an average S.D. 1.3% D. Exchange was quenched by mixing with cooled 1 M glycine-HCl, pH 2.3 at 1:1 ratio. A PAL DHR autosampler (CTC Analytics AG) controlled by Chronos software (AxelSemrau) was used to prepare HDX reactions and subsequent injection onto the LC system comprising a temperature-controlled box and Agilent Infinity II UPLC (Agilent Technologies) directly coupled to an ESI source of timsTOF Pro mass spectrometer (Bruker Daltonics). Here online proteolysis was done on a mixed bed pepsin/nepenthesin-2 column (bed volume 66 µL) and peptides were captured and desalted on the trap column (SecurityGuard™ ULTRA Cartridge UHPLC Fully Porous Polar C18, 2.1mm ID, Phenomenex) under the flow of 0.4% formic acid (FA) in water driven by the 1260 Infinity II Quaternary pump at the flow rate of 200 µL·min^-1^. Peptides were further separated on an analytical column (Luna Omega Polar C18, 1.6 µm, 100 Å, 1.0x100 mm, Phenomenex) using water-acetonitrile (ACN) gradient (10 – 45 % in 6 min; solvent A: 0.1% FA in water, solvent B: 0.1% FA, 2% water in ACN), which was delivered by the 1290 Infinity II LC pump under the flow rate of 40 µL·min^-1^. The LC outlet was directly interfaced ESI source of timsTOF Pro (Bruker Daltonics) mass spectrometer operated in the MS mode with a 1 Hz data acquisition. To minimize the loss of deuterium, the LC system was refrigerated to 0 °C. Fully deuterated control samples used for deuterium back-exchange correction were prepared for all variants [30].

Acquired LC-MS data were peak picked and exported in DataAnalysis (v. 5.3, Bruker Daltonics) and further analyzed by the DeutEx software [31]. Data visualization was performed using MSTools (http://peterslab.org/MSTools/index.php) [32]. For peptide identification, the same LC-MS system was used but the mass spectrometer acquired data in data-dependent MS/MS mode using PASEF. The LC-MS/MS data were searched against a customized database combining sequences of NQO1 (WT, Y126A/E/F and Y128A/E/F), common cRAP.fasta (https://www.thegpm.org/crap/) and proteases using MASCOT (v. 2.7, Matrix Science). Search parameters were set as follows: no enzyme, no modifications allowed, precursor tolerance 10 ppm, fragment ion tolerance 0.05 Da, decoy search enabled, FDR ˂ 1%, IonScore ˃ 20 and peptide length ˃ 5. The HDX dataset consisted of 295 peptides providing HDX data. The entire sequence of NQO1 was covered (100% sequence coverage) by peptides of an average peptide length of 11.2 residues and an average redundancy of 12.1.The HDX-MS data have been deposited to the ProteomeXchange Consortium via the PRIDE [33] partner repository with the dataset identifier PXD060224.

### 4.5 Pre-steady state enzyme kinetic analysis

Reductive processes for NQO1 variants upon HT from NAD(P)H were followed under anaerobic conditions using a stopped-flow spectrophotometer (SX.18MV, Applied Photophysics Ltd., Leatherhead, UK) interfaced with a photodiode array detector, essentially as described before [8]. Reactions were performed in 20 mM HEPES-KOH, pH 7.4. The flavin reductive half-reaction was measured by mixing NQO1 variants (7.5 µM) with NAD(P)H ranging from 7.5 to 100 µM. Multiple wavelength absorption data in the flavin absorption region were collected and processed as described [8]. Time-dependent spectral deconvolution was performed by global analysis and numerical integration methods using previously described procedures [8]. Deconvolution was carried out considering two sequential and irreversible steps (A→B→C), which correspond to the two catalytically relevant processes: A→B (*Fast process*) and B→C (*Slow process*) [8]. This procedure allowed to determine observed rate constants (*k*_obs_) for these steps, as well as spectroscopic properties of A, B and C species. When observed, hyperbolic dependences of *k*_obs_ vs. NADH concentrations were fitted using equation 1:

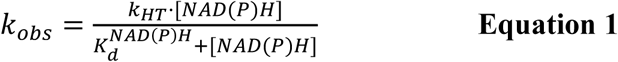

where *k*_HT_ is the limiting rate constant for HT and *K*d^NAD(P)H^ is the equilibrium dissociation constant for the coenzyme to a given active site.

### 4.6 Estimation of midpoint reduction potentials and photoreduction experiments

Reduction of NQO1 variants by the xanthine/XO method [21,34] was carried out in a closed anaerobic cuvette with a final concentration of ∼10 µM of each NQO1 variant, 2 µM benzyl viologen, 500 µM xanthine, 5 µM dye, 10 mM glucose and 10 U/mL glucose oxidase, in 20 mM HEPES-KOH pH 7.4. The indigo disulphonic acid two-electron exchanger (*E*_m_=-125 mV, *vs* standard hydrogen electrode) was the dye that better equilibrated with FAD reduction for all variants. After achieving anaerobic conditions, as above indicated using an anaerobic cuvette and a Schlenk line, 4 µg/mL of bovine milk XO (Sigma-Aldrich) were added from the side arm to the mixture, and spectra were recorded every 2 min scanning from 300 to 750 nm at 25 °C, in a CARY 3500 spectrophotometer (Agilent Technologies). To ensure thorough equilibration of the components during the experiment, the concentration of XO was set at 15 nM (within the recommended range of 1–100 nM, also replicated with 50 nM were also performed, providing the same values and therefore confirming equilibrium was reached) allowing the reductive process to proceed over a period of 2 to 3 h. Absorption changes accompanying reduction of the indicator dye and the flavoprotein at each point during the xanthine/XO reduction were used to determine the NQO1 *E*_ox/hq_ value using the Nernst equation as previously reported [21,34]. In short, being *E* of the flavin containing protein (P) and dye (D) as determined by their corresponding Nernst equations:

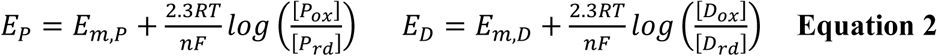

Slow rates of electron input by XO ensure the equilibrium of oxidized and reduced forms at any given time point of the process. Thus, the NQO1 and dye electrochemical potentials are assumed to be equal, and defining “*x*” and “*y*” as the Nernst concentration terms for the NQO1 and dye, respectively:

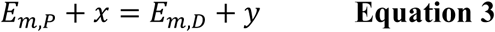

When the protein is in redox equilibrium [P_ox_] = [P_rd_], “*x* = 0” and “*y*” is defined as ΔE, the difference in midpoint potentials of NQO1 and the dye.

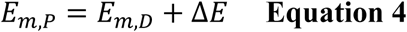

Thus, the difference in *E_m_* potential of NQO1 and the dye was calculated from a plot where the log([oxidized]/[reduced] ratio) for sample and dye of each spectrum is plotted against each other. In this particular case, the absorbance changes accompanying reduction of the indicator dye at 610 nm (where the NQO1 does not absorb) and of the flavoprotein at 468 nm (where the indicator has an isosbestic point) at each time point during the xanthine/XO reduction were used to calculate oxidized/reduced ratios for both the protein and the dye (equation 5).

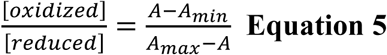

Where A is the observed absorbance with a given time point in the titration, and A_max_ and A_min_ the maximal and minimal absorbances observed. Plotting such “*x*” and “*y*” values on a graph and fitting them to equation 4 provides as intercept ΔE, the shift in midpoint potential between the FAD of the NQO1 variant and the dye.

Since experiments with WT NQO1 envisage transient potential stabilization of its anionic semiquinone, we also evaluated this feature by NQO1 (20 µM) in 20 mM HEPES-KOH pH 7.4, 3 mM EDTA, 4 µM 5-dRf (5-deazariboflavin), 5 mM glucose and 10 U/ml glucose oxidase in a closed anaerobic cuvette. The WT NQO1 sample was stepwise photoirradiated for 5 s periods with a blue LED strip tube that provides an intensity of 900 µmol photons/(m^2^·s) around the cuvette. Photoreduction was followed by recording the visible spectra after each irradiation by using a CARY 3500 spectrophotometer (Agilent Technologies).

## Author Contributions

P.M., M.M. and A.L.P. planned experiments; J.L.P-G, M.R., P.V. and D.S.L. performed experiments; All authors analyzed data; P.M., M.M. and A.L.P. contributed reagents or other essential material; M.M. and A.L.P drafted the paper; All authors contributed to the final version of the paper.

## Acknowledgements

A.L.P. acknowledges funding from ERDF/Spanish Ministry of Science, Innovation and Universities—State Research Agency (Grant number RTI2018-096246-B-I00), Consejería de Economía, Conocimiento, Empresas y Universidad, Junta de Andalucía (Grant number P18-RT-2413), ERDF/Counseling of Economic transformation, Industry, Knowledge and Universities (Grant B-BIO-84-UGR20). M.M. acknowledges funding from the Spanish State Research Agency and FEDER (MCIN/AEI-FEDER, Grant PID2022-136369NB-I00) and the Government of Aragón-FEDER (Grant number E35_23R). P.M. acknowledges support from the MEYS project OP JAK INTER-MICRO (CZ.02.01.01/00/22_008/0004597). Access to the Instruct-CZ center (BioCeV), was supported by CIISB (LM2023042 and CZ.02.01.01/00/23_015/0008175) and Instruct- Internship to J.L.P-G. (PID 2545).

## References

1 Salido E, Timson DJ, Betancor-Fernández I, Palomino-Morales R, Anoz-Carbonell E, Pacheco-García JL, Medina M & Pey AL (2022) Targeting HIF-1α Function in Cancer through the Chaperone Action of NQO1: Implications of Genetic Diversity of NQO1. J Pers Med 12, 747.

2 Ross D & Siegel D (2021) The diverse functionality of NQO1 and its roles in redox control. Redox Biol 41, 101950.

3 Asher G, Tsvetkov P, Kahana C & Shaul Y (2005) A mechanism of ubiquitin-independent proteasomal degradation of the tumor suppressors p53 and p73. Genes Dev 19, 316–21.

4 Beaver SK, Mesa-Torres N, Pey AL & Timson DJ (2019) NQO1: A target for the treatment of cancer and neurological diseases, and a model to understand loss of function disease mechanisms. Biochim Biophys Acta Proteins Proteom 1867, 663–676.

5 Asher G, Dym O, Tsvetkov P, Adler J & Shaul Y (2006) The crystal structure of NAD(P)H quinone oxidoreductase 1 in complex with its potent inhibitor dicoumarol. Biochemistry 45, 6372–8.

6 Grieco A, Boneta S, Gavira JA, Pey AL, Basu S, Orlans J, Sanctis D de, Medina M & Martin-Garcia JM (2024) Structural dynamics and functional cooperativity of human NQO1 by ambient temperature serial crystallography and simulations. Protein Sci 33, e4957.

7 Medina-Carmona E, Rizzuti B, Martín-Escolano R, Pacheco-García JL, Mesa-Torres N, Neira JL, Guzzi R & Pey AL (2019) Phosphorylation compromises FAD binding and intracellular stability of wild-type and cancer-associated NQO1: Insights into flavo-proteome stability. Int J Biol Macromol 125, 1275–1288.

8 Anoz-Carbonell E, Timson DJ, Pey AL & Medina M (2020) The catalytic cycle of the antioxidant and cancer-associated human NQO1 enzyme: Hydride transfer, conformational dynamics and functional cooperativity. Antioxidants 9, 1–22.

9 Pacheco-Garcia JL, Anoz-Carbonell E, Vankova P, Kannan A, Palomino-Morales R, Mesa-Torres N, Salido E, Man P, Medina M, Naganathan AN & Pey AL (2021) Structural basis of the pleiotropic and specific phenotypic consequences of missense mutations in the multifunctional NAD(P)H:quinone oxidoreductase 1 and their pharmacological rescue. Redox Biol 46, 102112.

10 Megarity CF, Abdel-Aal Bettley H, Caraher MC, Scott KA, Whitehead RC, Jowitt TA, Gutierrez A, Bryce RA, Nolan KA, Stratford IJ & Timson DJ (2019) Negative Cooperativity in NAD(P)H Quinone Oxidoreductase 1 (NQO1). Chembiochem 20, 2841–2849.

11 Clavería-Gimeno R, Velazquez-Campoy A & Pey AL (2017) Thermodynamics of cooperative binding of FAD to human NQO1: Implications to understanding cofactor-dependent function and stability of the flavoproteome. Arch Biochem Biophys 636, 17–27.

12 Pey AL, Megarity CF & Timson DJ (2014) FAD binding overcomes defects in activity and stability displayed by cancer-associated variants of human NQO1. Biochim Biophys Acta Mol Basis Dis 1842, 2163–2173

13 Betancor-Fernández I, Timson DJ, Salido E & Pey AL (2018) Natural (and Unnatural) Small Molecules as Pharmacological Chaperones and Inhibitors in Cancer. Handb Exp Pharmacol 245, 155–190.

14 Islam F, Basilone N, Yoo V, Ball E & Shilton B (2024) Evolutionary analysis of Quinone Reductases 1 and 2 suggests that NQO2 evolved to function as a pseudoenzyme. Protein Sci 33, e5234.

15 Vankova P, Salido E, Timson DJ, Man P & Pey AL (2019) A Dynamic Core in Human NQO1 Controls the Functional and Stability Effects of Ligand Binding and Their Communication across the Enzyme Dimer. Biomolecules 9, 728.

16 Pacheco-Garcia JL, Anoz-Carbonell E, Loginov DS, Vankova P, Salido E, Man P, Medina M, Palomino-Morales R & Pey AL (2022) Different phenotypic outcome due to site-specific phosphorylation in the cancer-associated NQO1 enzyme studied by phosphomimetic mutations. Arch Biochem Biophys 729, 109392.

17 Pacheco-Garcia JL, Loginov DS, Anoz-Carbonell E, Vankova P, Palomino-Morales R, Salido E, Man P, Medina M, Naganathan AN & Pey AL (2022) Allosteric Communication in the Multifunctional and Redox NQO1 Protein Studied by Cavity-Making Mutations. Antioxidants 11, 1110.

18 Megarity CF & Timson DJ (2019) Cancer-associated variants of human NQO1: impacts on inhibitor binding and cooperativity. Biosci Rep 39, BSR20191874.

19 Medina-Carmona E, Palomino-Morales RJ, Fuchs JE, Padín-Gonzalez E, Mesa-Torres N, Salido E, Timson DJ & Pey AL (2016) Conformational dynamics is key to understanding loss-of-function of NQO1 cancer-associated polymorphisms and its correction by pharmacological ligands. Sci Rep 6, 20331.

20 Pacheco-García JL, Anoz-Carbonell E, Loginov DS, Kavan D, Salido E, Man P, Medina M & Pey AL (2023) Counterintuitive structural and functional effects due to naturally occurring mutations targeting the active site of the disease-associated NQO1 enzyme. FEBS J 290, 1855–1873.

21 Maklashina E & Cecchini G (2020) Determination of Flavin Potential in Proteins by Xanthine/Xanthine Oxidase Method. Bio Protoc 10, e3571.

22 Tedeschi G, Chen S & Massey V (1995) DT-diaphorase. Redox potential, steady-state, and rapid reaction studies. J Biol Chem 270, 1198–204.

23 Hornbeck P V, Zhang B, Murray B, Kornhauser JM, Latham V & Skrzypek E (2015) PhosphoSitePlus, 2014: mutations, PTMs and recalibrations. Nucleic Acids Res 43, D512-20.

24 Pacheco-García JL, Cano-Muñoz M, Sánchez-Ramos I, Salido E & Pey AL (2020) Naturally-Occurring Rare Mutations Cause Mild to Catastrophic Effects in the Multifunctional and Cancer-Associated NQO1 Protein. J Pers Med 10, 207.

25 Pacheco-Garcia JL, Cagiada M, Tienne-Matos K, Salido E, Lindorff-Larsen K & L. Pey A (2022) Effect of naturally-occurring mutations on the stability and function of cancer-associated NQO1: Comparison of experiments and computation. Front Mol Biosci 9, 1063620.

26 Lienhart W-D, Gudipati V, Uhl MK, Binter A, Pulido SA, Saf R, Zangger K, Gruber K & Macheroux P (2014) Collapse of the native structure caused by a single amino acid exchange in human NAD(P)H:quinone oxidoreductase(1.). FEBS J 281, 4691– 4704.

27 Lienhart W-D, Strandback E, Gudipati V, Koch K, Binter A, Uhl MK, Rantasa DM, Bourgeois B, Madl T, Zangger K, Gruber K & Macheroux P (2017) Catalytic competence, structure and stability of the cancer-associated R139W variant of the human NAD(P)H:quinone oxidoreductase 1 (NQO1). FEBS J 284, 1233–1245.

28 Medina-Carmona E, Fuchs JE, Gavira JA, Mesa-Torres N, Neira JL, Salido E, Palomino-Morales R, Burgos M, Timson DJ & Pey AL (2017) Enhanced vulnerability of human proteins towards disease-associated inactivation through divergent evolution. Hum Mol Genet 26, 3531–3544.

29 Pacheco-Garcia JL, Loginov D, Rizzuti B, Vankova P, Neira JL, Kavan D, Mesa-Torres N, Guzzi R, Man P & Pey AL (2022) A single evolutionarily divergent mutation determines the different FAD-binding affinities of human and rat NQO1 due to site-specific phosphorylation. FEBS Lett 596, 29–41.

30 Vankova P, Pacheco-Garcia JL, Loginov DS, Gómez-Mulas A, Kádek A, Martín- Garcia JM, Salido E, Man P & Pey AL (2024) Insights into the pathogenesis of primary hyperoxaluria type I from the structural dynamics of alanine:glyoxylate aminotransferase variants. FEBS Lett 598, 485–499.

31 Trcka F, Durech M, Vankova P, Chmelik J, Martinkova V, Hausner J, Kadek A, Marcoux J, Klumpler T, Vojtesek B, Muller P & Man P (2019) Human Stress-inducible Hsp70 Has a High Propensity to Form ATP-dependent Antiparallel Dimers That Are Differentially Regulated by Cochaperone Binding*. Molecular & Cellular Proteomics 18, 320–337.

32 Kavan D & Man P (2011) MSTools—Web based application for visualization and presentation of HXMS data. Int J Mass Spectrom 302, 53–58.

33 Perez-Riverol Y, Bai J, Bandla C, García-Seisdedos D, Hewapathirana S, Kamatchinathan S, Kundu DJ, Prakash A, Frericks-Zipper A, Eisenacher M, Walzer M, Wang S, Brazma A & Vizcaíno JA (2022) The PRIDE database resources in 2022: a hub for mass spectrometry-based proteomics evidences. Nucleic Acids Res 50, D543–D552.

34 Christgen SL, Becker SM & Becker DF (2019) Methods for determining the reduction potentials of flavin enzymes. Methods Enzymol 620, 1–25.

