## Supplementary information for "Active site tyrosine residues in human NQO1 homodimer are critical for non-synchronous enzyme catalysis at the two active sites"

**Figure S1. SDS-PAGE (12% acrylamide) analysis of purified NQO1 variants.** 10 μg of each variant were loaded per lane.

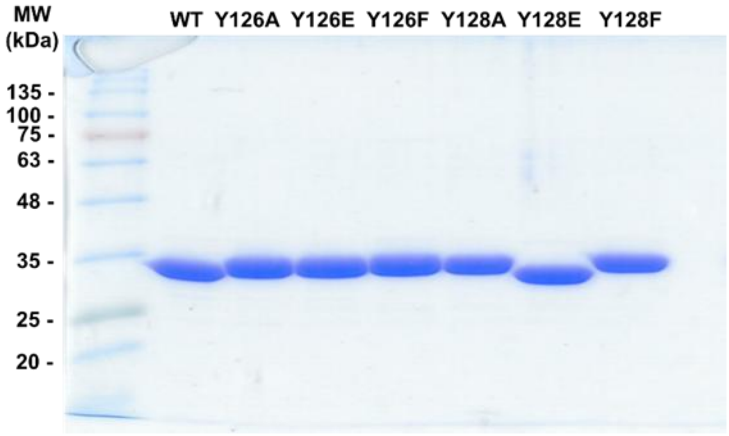


**Figure S2. UV-visible spectra of NQO1 variants as purified.** (A) Spectra recorded at ~20 μM in NQO1 protein monomer. (B) FAD content of purified variants determined from spectra in panel A according to [1]. Data are the average from two independent purifications of each variant and thus errors are not displayed.

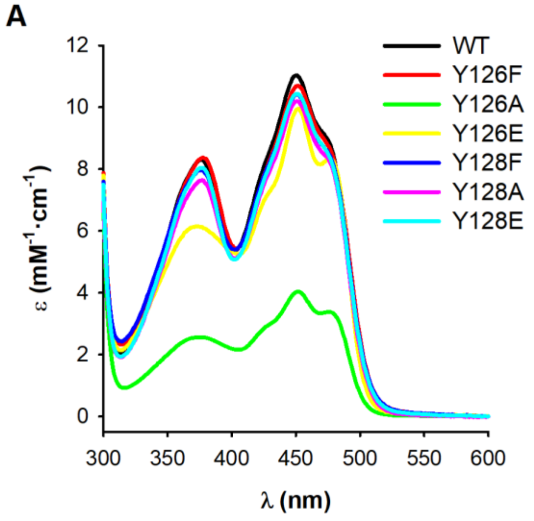


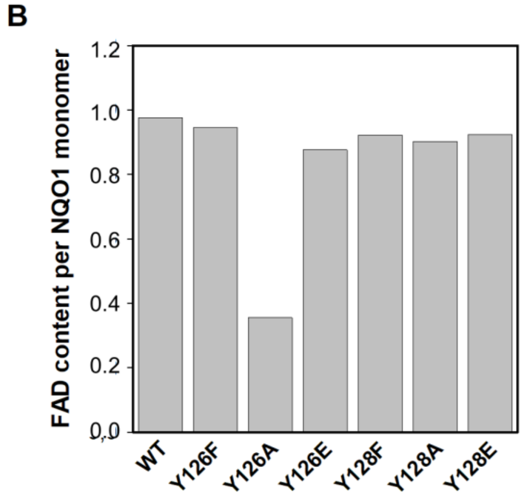

**Figure S3. Thermal denaturation of NQO1 by fluorescence spectroscopy.** (A) Normalized thermal denaturation profiles; Data for a single representative experiment for each variant is shown for the sake of clarity. (B) $T_m$ values determined for thermal denaturation of NQO1 variants (mean±s.d. from three replicates).

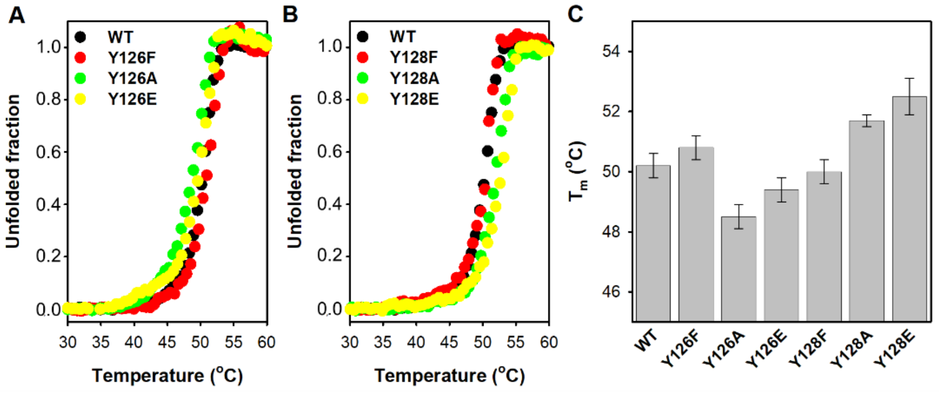

**Figure S4. HDX of WT NQO1 in the absence (black symbols) or the presence of an excess of $NAD^+$ (red symbols).** Lines are best fits to $D(t)= D_\infty+A\cdot e^{(-k_{obs}\cdot t)}$, where D(t) is the time dependent % of deuteration, $D_\infty$ is the % of deuteration at $t\rightarrow\infty$, A is the amplitude of observed first-order phase (% of exchange) and $k_{obs}$ is the first-order rate constant for this phase. Data correspond to segments comprising residues 1-42 (excluding Met1).

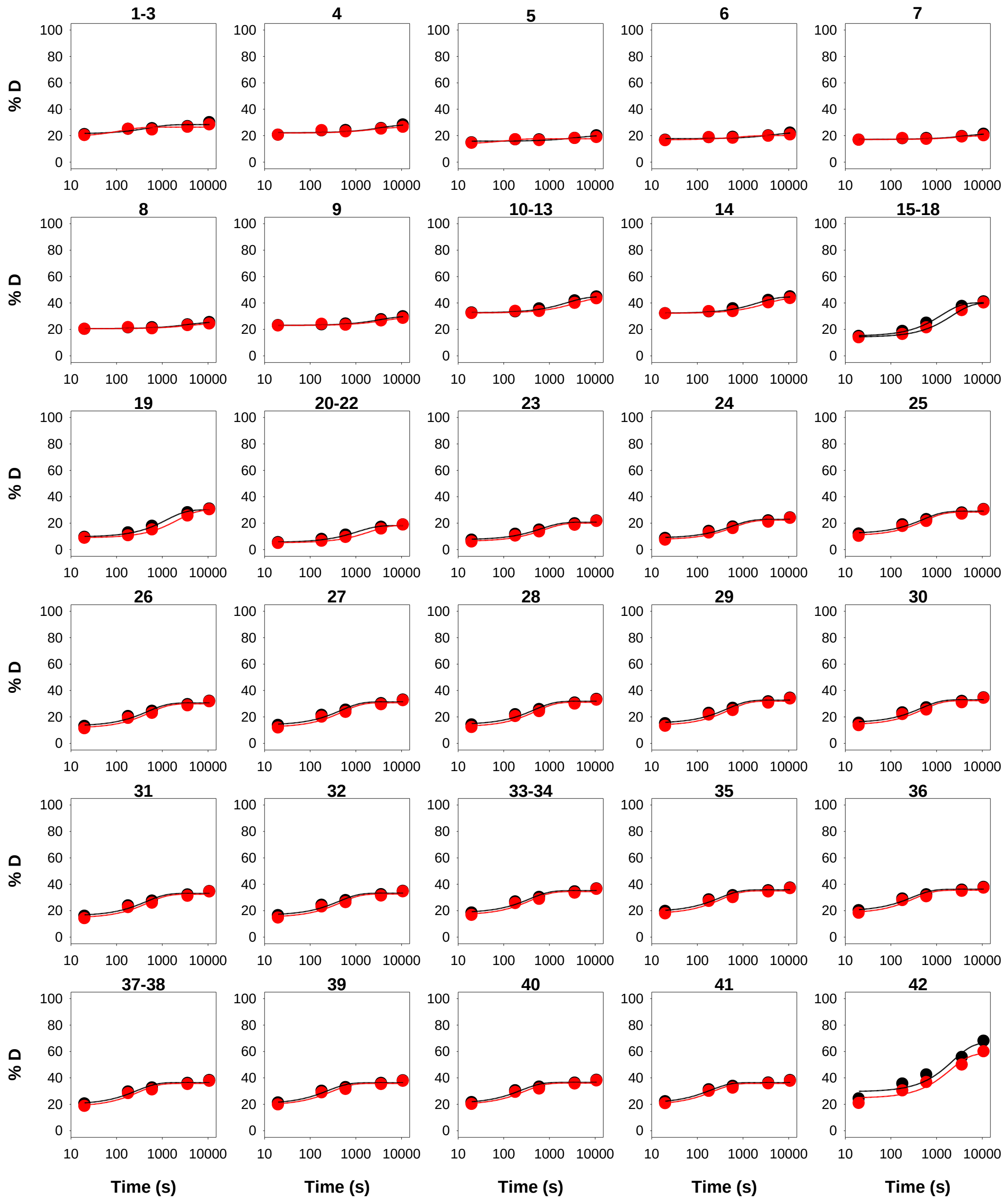

**Figure S5. HDX of WT NQO1 in the absence (black symbols) or the presence of an excess of $NAD^+$ (red symbols).** Lines are best fits to $D(t)= D_\infty+A\cdot e^{(-k_{obs}\cdot t)}$, where D(t) is the time dependent % of deuteration, $D_\infty$ is the % of deuteration at $t\rightarrow\infty$, A is the amplitude of observed first-order phase (% of exchange) and $k_{obs}$ is the first-order rate constant for this phase. Data correspond to segments comprising residues 43-97 (excluding Met1).

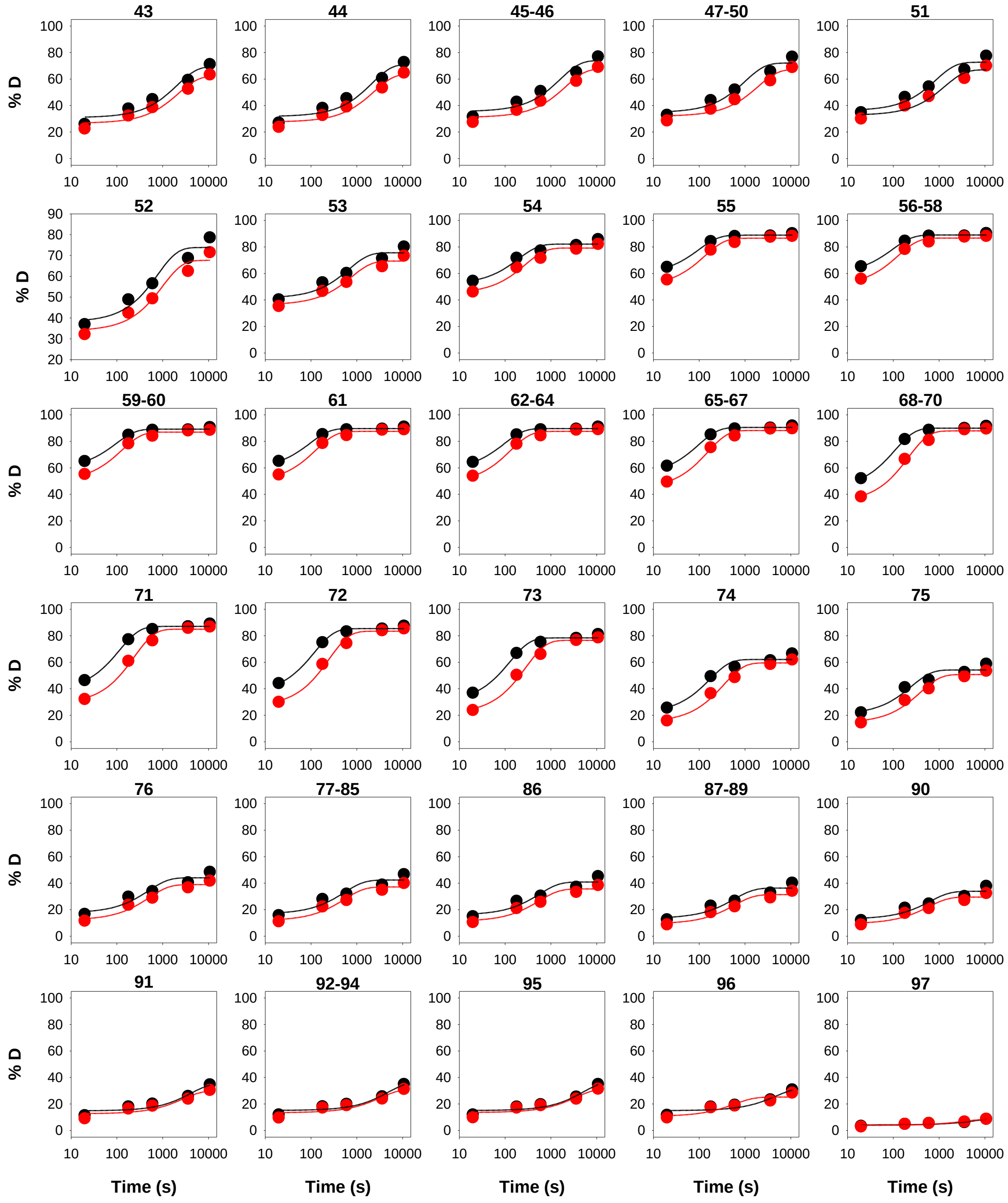

**Figure S6. HDX of WT NQO1 in the absence (black symbols) or the presence of an excess of NAD$^+$ (red symbols).** Lines are best fits to $D(t)= D_{\infty}+A\cdot e^{(-k_{obs}\cdot t)}$, where D(t) is the time dependent % of deuteration, $D_{\infty}$ is the % of deuteration at $t\rightarrow\infty$, A is the amplitude of observed first-order phase (% of exchange) and $k_{obs}$ is the first-order rate constant for this phase. Data correspond to segments comprising residues 98-164 (excluding Met1).

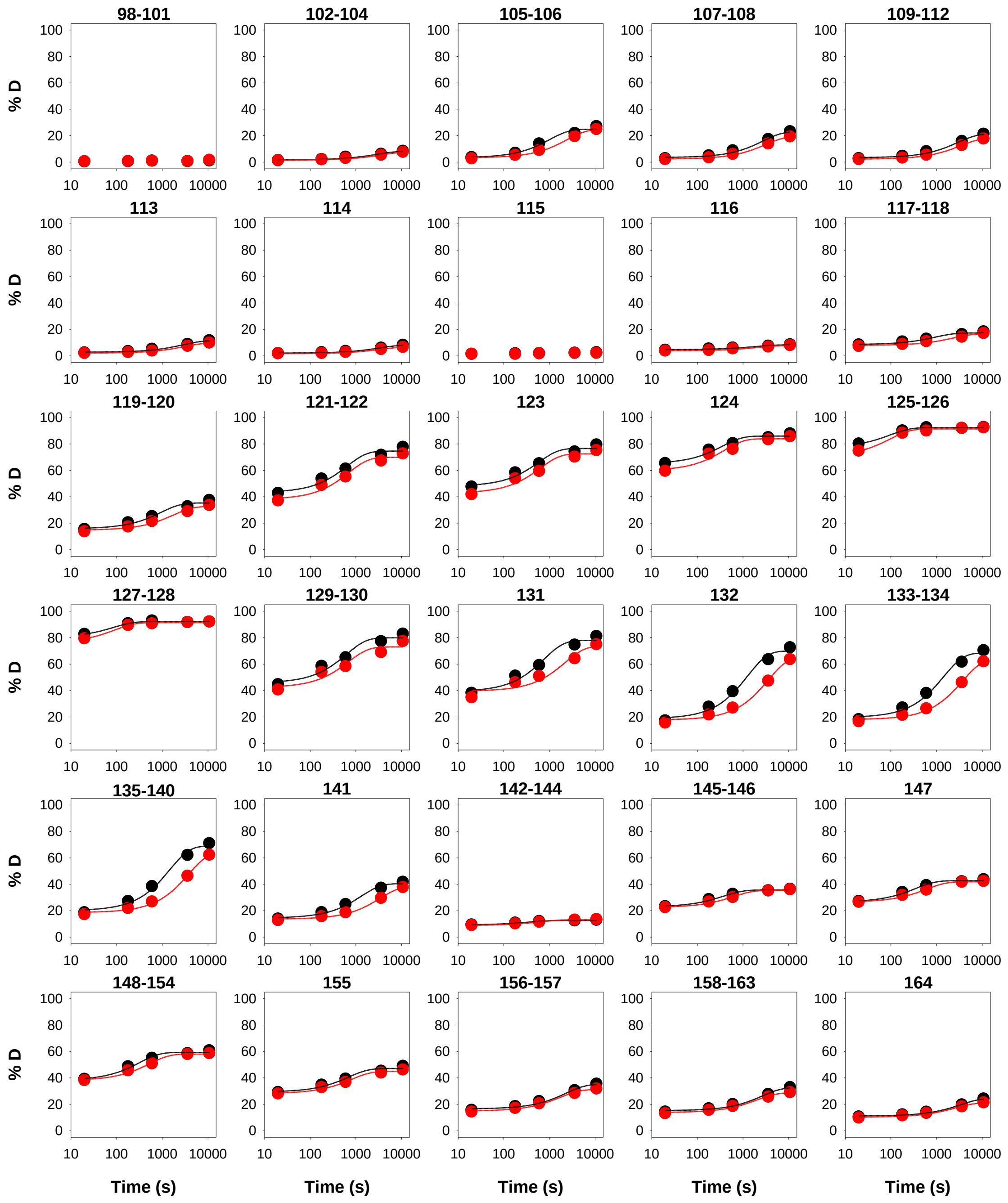

**Figure S7. HDX of WT NQO1 in the absence (black symbols) or the presence of an excess of $NAD^+$ (red symbols).** Lines are best fits to $D(t)= D_\infty+A\cdot e^{(-k_{obs}\cdot t)}$, where D(t) is the time dependent % of deuteration, $D_\infty$ is the % of deuteration at $t\rightarrow\infty$, A is the amplitude of observed first-order phase (% of exchange) and $k_{obs}$ is the first-order rate constant for this phase. Data correspond to segments comprising residues 165-211 (excluding Met1).

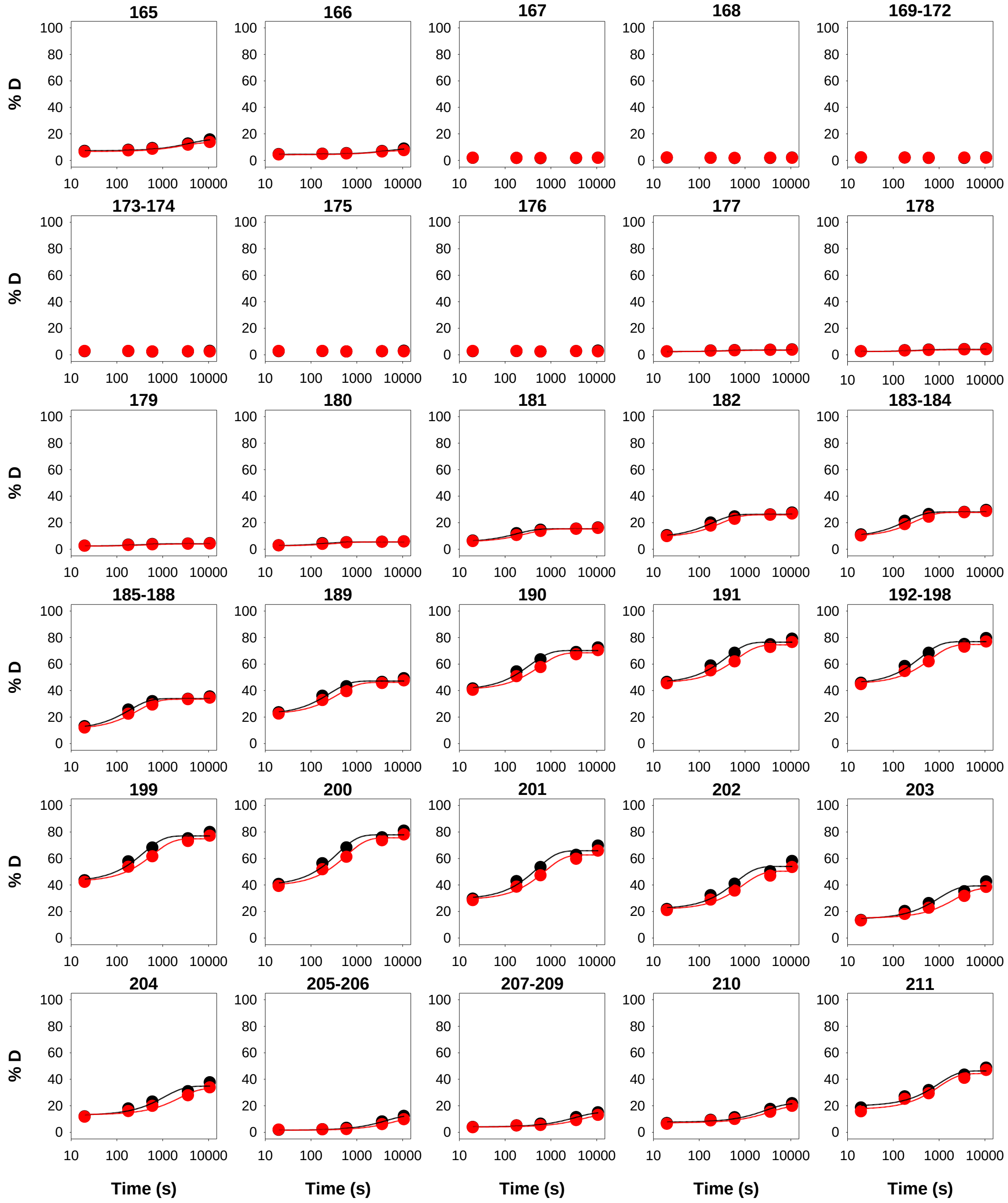

**Figure S8. HDX of WT NQO1 in the absence (black symbols) or the presence of an excess of $NAD^+$ (red symbols).** Lines are best fits to $D(t)= D_\infty+A\cdot e^{(-k_{obs}\cdot t)}$, where D(t) is the time dependent % of deuteration, $D_\infty$ is the % of deuteration at $t\rightarrow\infty$, A is the amplitude of observed first-order phase (% of exchange) and $k_{obs}$ is the first-order rate constant for this phase. Data correspond to segments comprising residues 212-273 (excluding Met1).

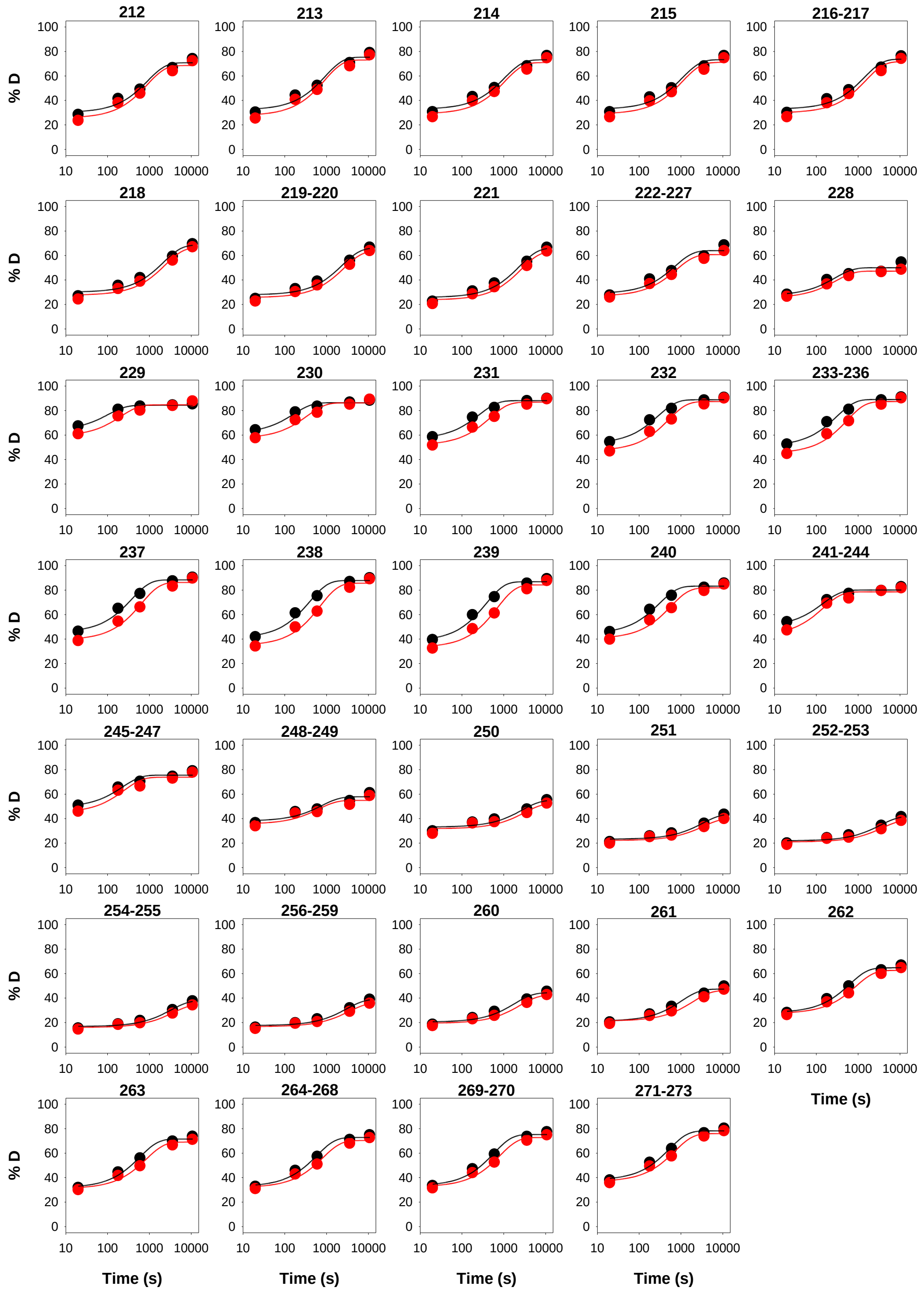

**Figure S9. Effect of NAD$^+$ on the HDX of the different NQO1 variants.** (A) WT and Y126 and (B) WT and Y128. The variants Y126F and Y128F are not displayed due to their marginal effects (see Figure 2 in the main text). The plots show the difference between each variant in the presence or absence of NAD$^+$ determined as described in [2].

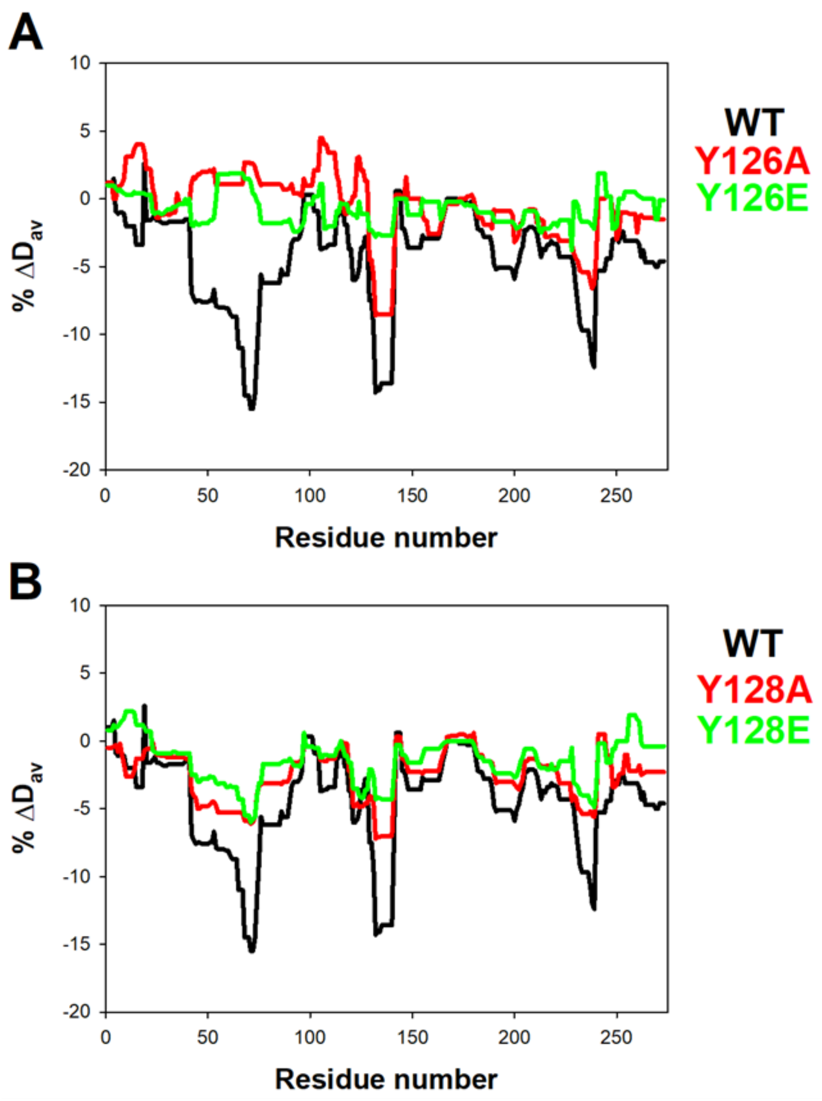

**Figure S10. SDS-PAGE (12% acrylamide) analysis of NQO1 variants degraded by thermolysin.** The NQO1 variant, concentration of thermolysin used and incubation time is indicated in each panel. Molecular weight markers are also shown at the left side of the gels.

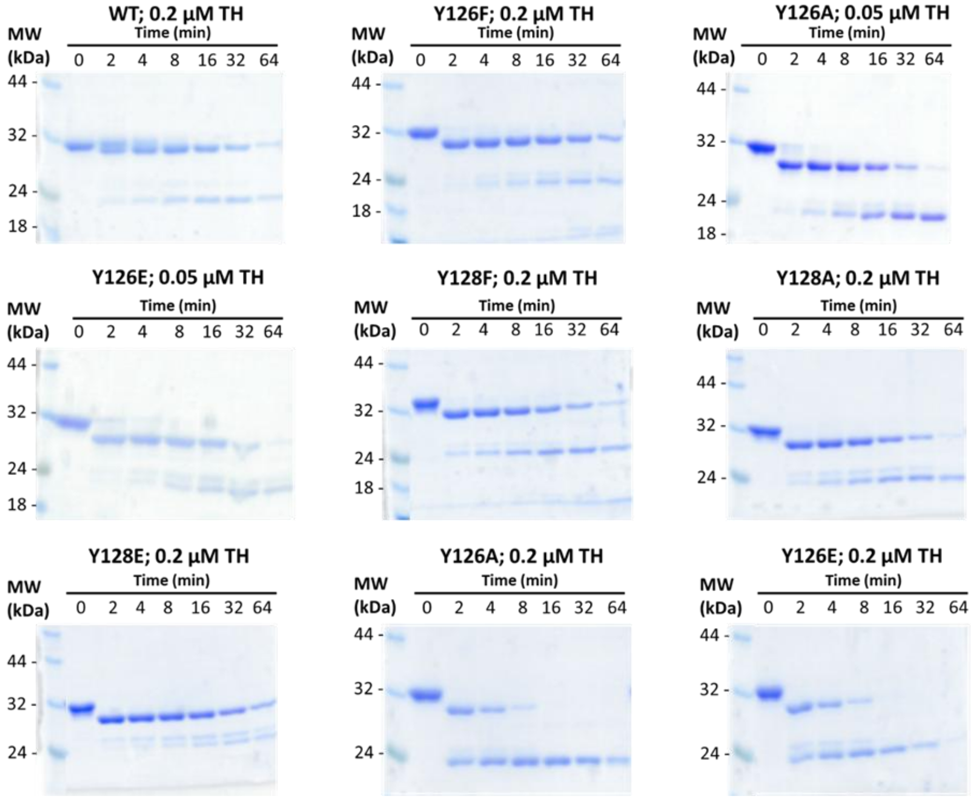

**Figure S11. Time-dependent spectra of the flavin reduction of $NQO1_{ox}$ variants by NADH**. Spectral evolution after mixing NQO1 (7.5 μM) with NADH (20 μM) in 20 mM HEPES-KOH, pH 7.4, at 6 °C. Panels A-F correspond to the indicated variants. Different coloured lines show the spectra at different reaction times as indicated in the corresponding legends. The dark grey line represents the spectrum of NQO1 before mixing with NADH. Absorption is shown in AU. Data are from representative from n > 3.

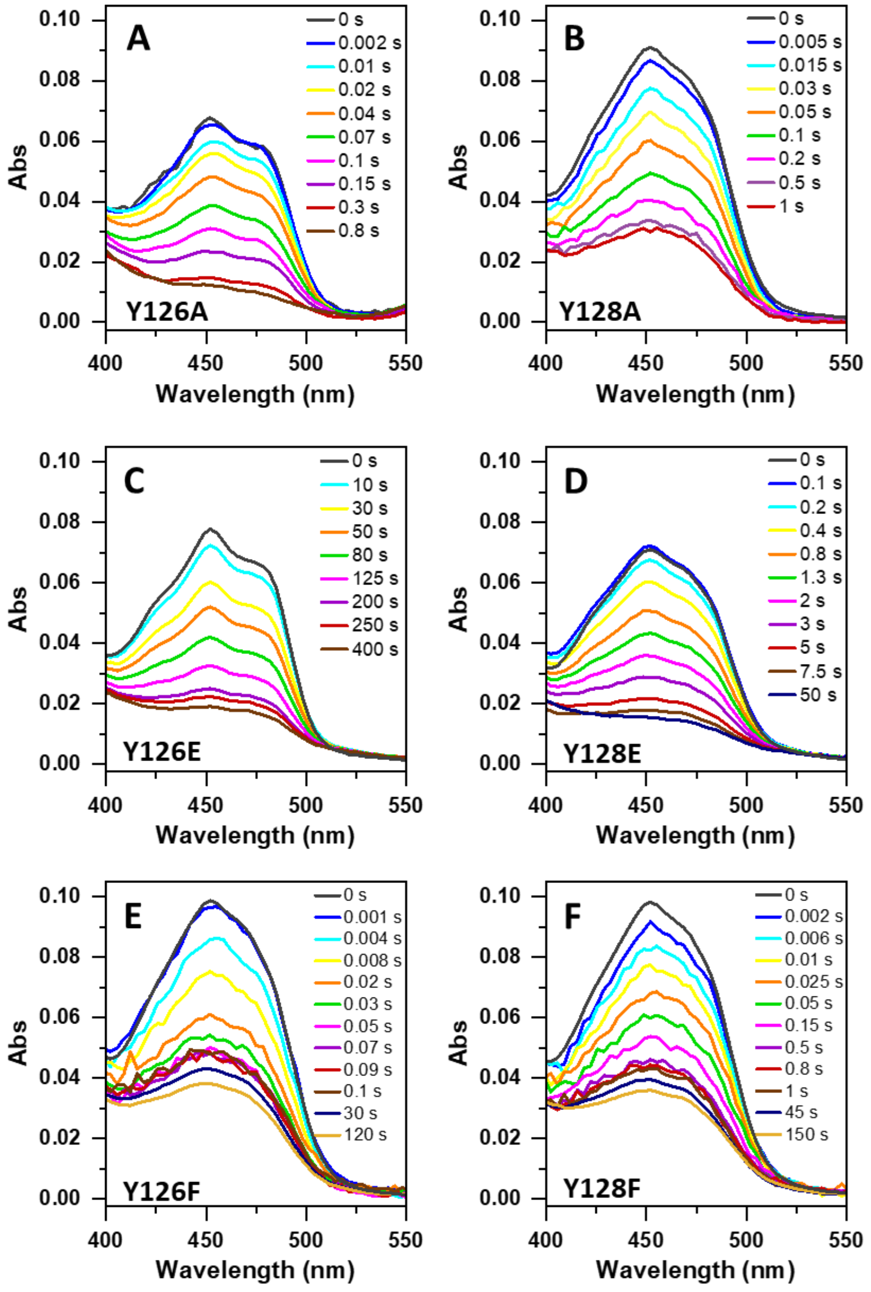

**Figure S12. Evaluation of individual kinetic processes for the flavin reduction of NQO1 variants by NADH**. A-F showed on the left panels the deconvolution into spectral species along the process for the indicated NQO1 variants. The % of the total $A_{450}$ evolved during the entire reaction is indicated, as well as the % of the $A_{450nm}$ for the overall reaction corresponding to the *fast* (A→B, species A is transformed into species B) and *slow* (B→C) steps. In the case of Y126F and Y128F variants, an additional (C→D) step describes a final very slow process that is observed separated from the major ones by a lag phase. A-F show on the right panels time evolution for flavin reduction as followed by changes in $A_{450nm}$ (black open circles). Lines are best fits to a two-step model (A→B→C) (green lines). For Y126F and Y128F mutants a lag phase separates those processes from a final very slow process that accounts for a very small amplitude change: in these cases, this step (C→D) has been independently fitted. The lower right panels show the corresponding residuals (black open circles) for data fitting. $A_{450nm}$ and $\Delta A_{450nm}$ are shown in mAU. In all cases, shown data correspond to representative transients recorded after mixing NQO1 (7.5 µM) with NADH (20 µM) in 20 mM HEPES-KOH, pH 7.4, at 6 °C (raw data presented in Figure S11). Data are representative from n > 3.

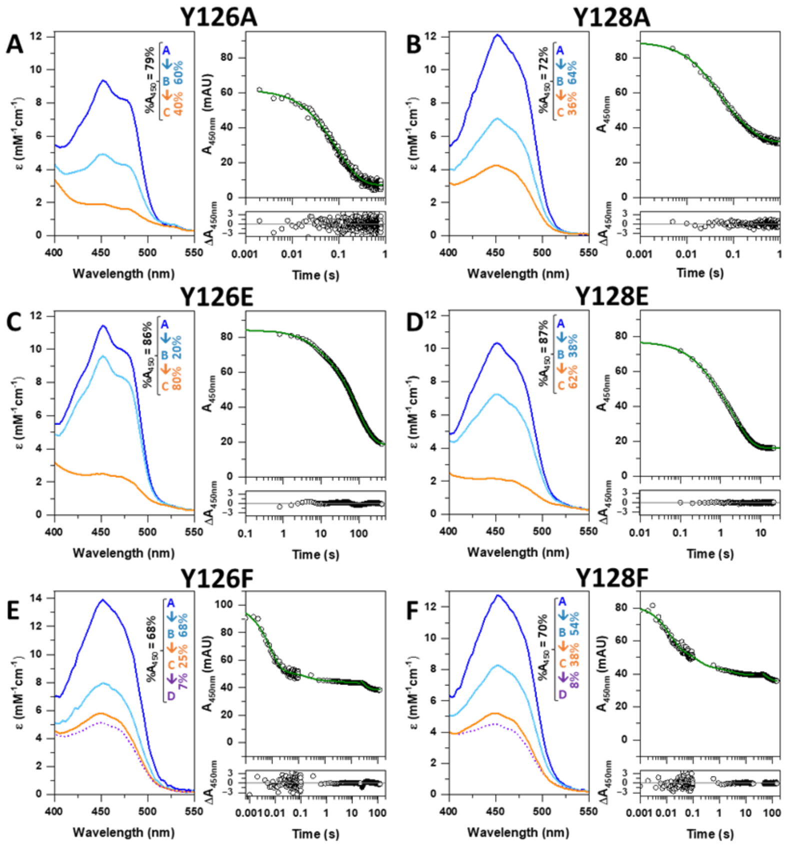

**Figure S13. Time-dependent spectra of the flavin reduction of NQO1$_{ox}$ variants by NADPH**. Spectral evolution after mixing NQO1 (7.5 µM) with NADPH (20 µM) in 20 mM HEPES-KOH, pH 7.4, at 6 °C. Panels A-F correspond to the indicated variants. Different coloured lines show the spectra at different reaction times as indicated in the corresponding legends. The dark grey line represents the spectrum of NQO1$_{ox}$ before mixing. Absorption is shown in AU. Data are from a single measurement and representative from n > 3.

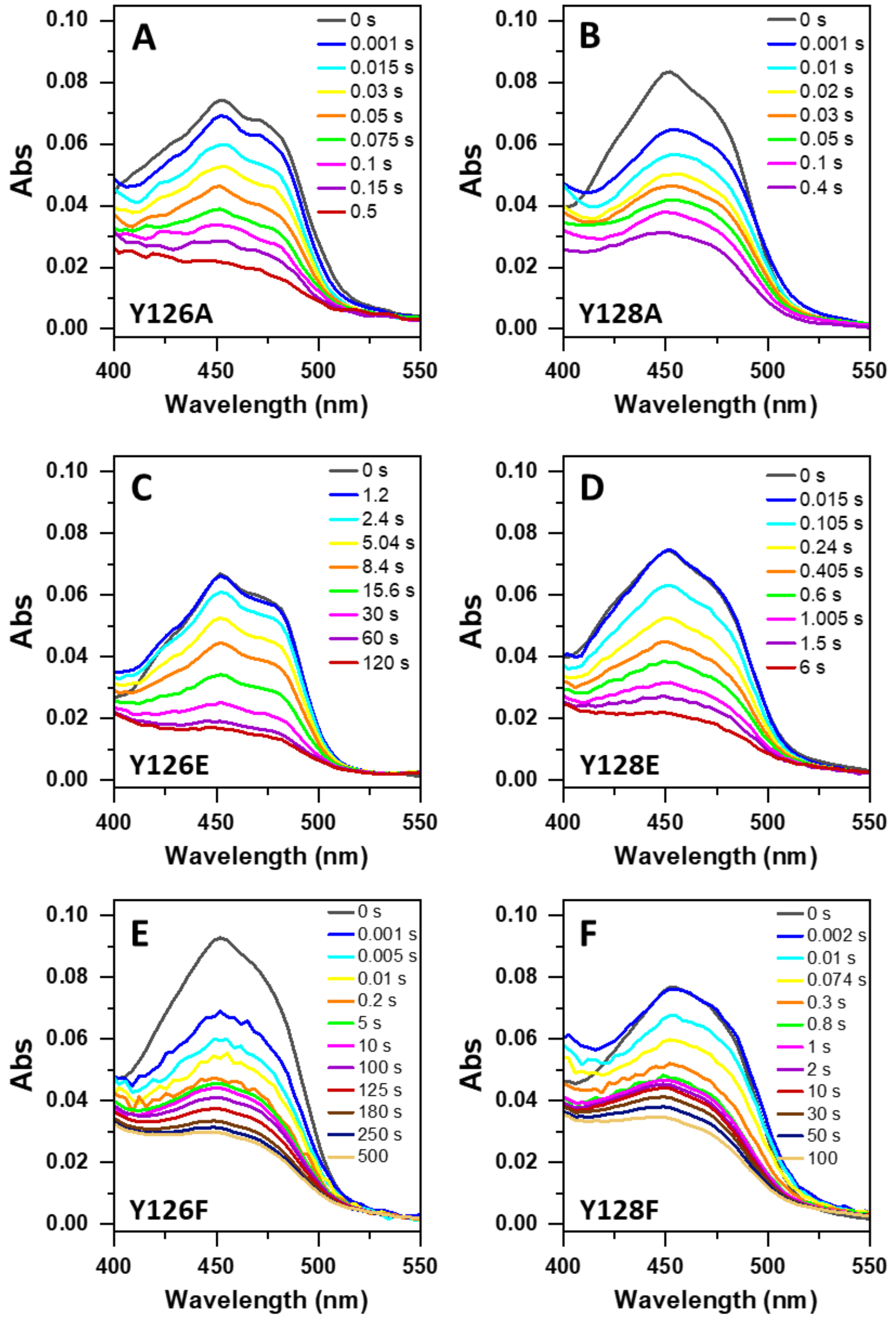

**Figure S14. Evaluation of individual kinetic processes for the flavin reduction of NQO1 variants by NADPH**. A-F show on the left panels the deconvolution in to spectral species along the process for the indicated NQO1 variants. The % of the total A450 evolved during the entire reaction is indicated, as well as the % of the $A_{450nm}$ for the overall reaction corresponding to the *fast* (A→B, species A is transformed into species B) and *slow* (B→C) steps. In the case of Y126F and Y128F variants, and additional (C→D) step describes a final very slow process that is observed separated from the major one by a lag phase. A-F show on the right panels time evolution for flavin reduction as followed by changes in $A_{450nm}$ (black open circles). Lines are best fits to a two-step model (A→B→C) (green lines). For Y126F and Y128F mutants a lag phase separates those processes from a final very slow process that accounts for a very small amplitude change: in these cases, this step (C→D) has been independently fitted. The lower right panels show the corresponding residuals (black open circles) for data fitting. $A_{450nm}$ and $\Delta A_{450nm}$ are shown in mAU. In all cases, shown data correspond to representative transients recorded after mixing $NQO1_{ox}$ (7.5 µM) with NADPH (20 µM) in 20 mM HEPES-KOH, pH 7.4, at 6 °C (raw data presented in Figure S13). Data are representative from n > 3.

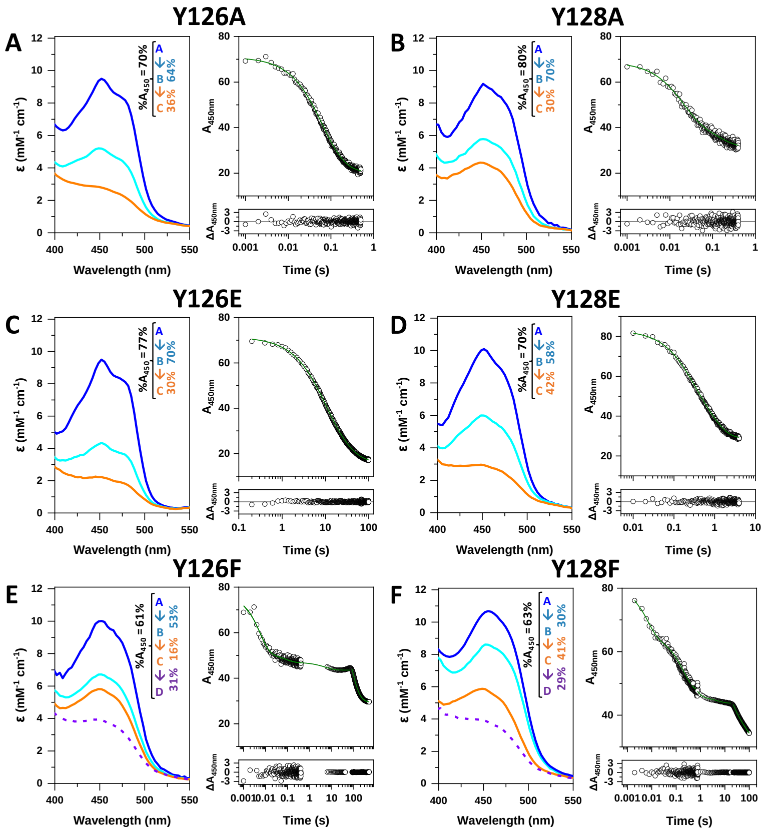

**Figure S15. Midpoint reduction potential of NQO1 variants.** (A) Spectral evolution upon mid-point reduction potential assessment of WT NQO1 using the xanthine/XO system and indigo disulphonic acid as a dye. Coloured Vis absorption spectra from blue to orange degradation correspond to time points evolution along protein reduction. Spectra were recorded every 2 min for up to 3 h in 20 mM HEPES-KOH, pH 7.4, at 25 °C. (B) Logarithm of the ratios of oxidized/reduced NQO1 and oxidized/reduced dye at different time points for the different variants when evaluated as in (A). Spectral evolution during (C) photoreduction of WT NQO1: spectra were recorded at different stepwise during light irradiation in 20 mM HEPES-KOH, pH 7.4, 3 mM EDTA, 4 μM 5-dRf at 25 °C; and (D) reoxidation of WT NQO1: spectra recorded stepwise upon opening of the cuvette in (C) to the air. Coloured Vis absorption spectra from blue to orange degradation correspond to different points along protein reduction/reoxidation. Experiments were reproduced at least twice.

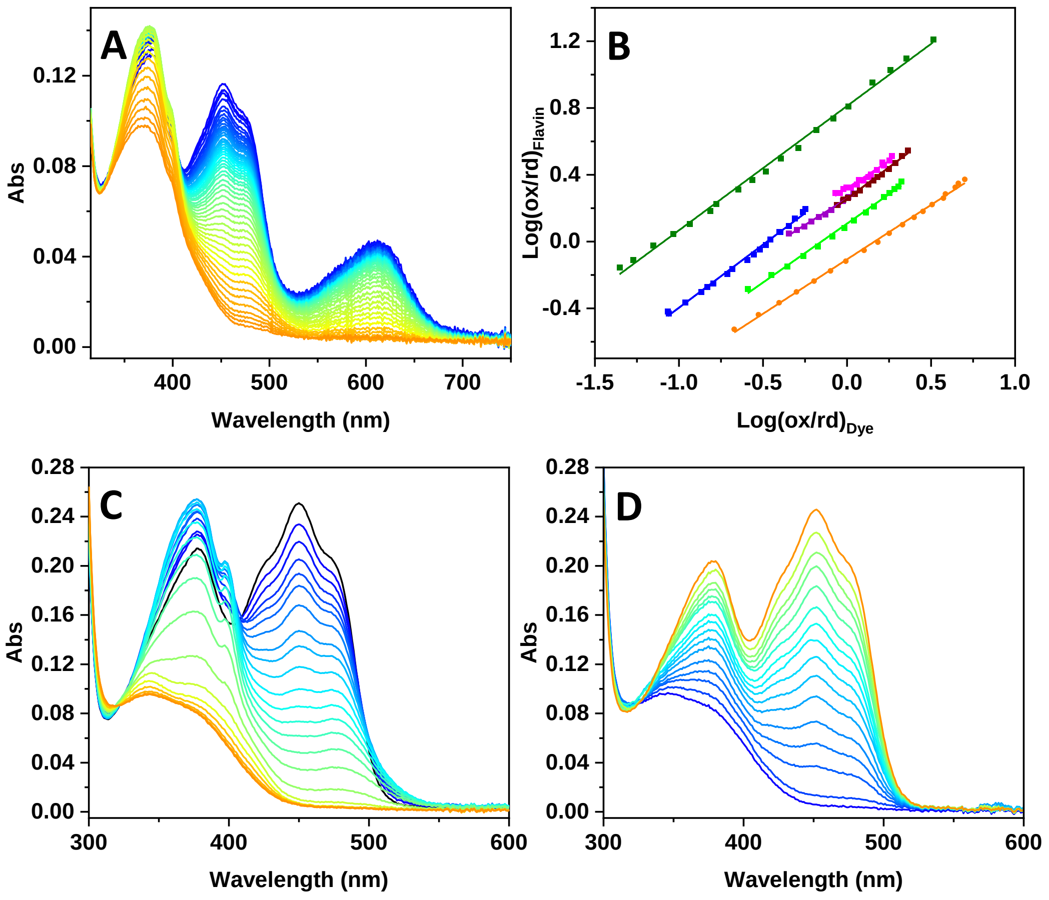

**Table S1. Best-fit parameters for the HDX of NQO1 in the absence or presence of $NAD^+$ using an exponential three-parameters fitting function as follows: $D(t)= D_\infty+A\cdot e^{(-k_{obs}\cdot t)}$.** Errors are those from fittings, and those for $D_0$ were linearly propagated. N.Det.- not determined, because the HDX was too low.

| Segment | $NAD^+$ | $D_\infty$ (%) | $D_0$ (%) | A (%) | $k_{obs}$ ($s^{-1}$) | $r^2$ |
|---|---|---|---|---|---|---|
| **1-3** | - | 28.4±1.4 | 18.5±3.2 | 6.9±2.4 | $2.8±1.8\cdot10^{-3}$ | 0.814 |
| | + | 26.5±1.2 | 19.2±3.4 | 7.3±2.9 | $8.8±8.1\cdot10^{-3}$ | 0.791 |
| **4** | - | 28.3±2.6 | 22.2±3.6 | 6.1±2.6 | $2.8±1.8\cdot10^{-4}$ | 0.818 |
| | + | 26.4±1.8 | 22.0±2.5 | 4.4±1.9 | $4.2±6.1\cdot10^{-4}$ | 0.864 |
| **5** | - | 20.2±2.1 | 15.8±2.5 | 4.4±2.0 | $2.4±3.2\cdot10^{-4}$ | 0.823 |
| | + | 17.7±0.7 | 13.6±2.3 | 4.1±1.8 | $8.9±10.1\cdot10^{-3}$ | 0.764 |
| **6** | - | 22.5±2.3 | 17.7±3.0 | 4.8±2.2 | $2.1±2.5\cdot10^{-4}$ | 0.866 |
| | + | 20.1±0.8 | 16.9±1.8 | 3.2±1.2 | $1.3±1.5\cdot10^{-3}$ | 0.781 |
| **7** | - | 21.6±1.0 | 17.2±2.5 | 4.4±1.0 | $2.2±1.3\cdot10^{-4}$ | 0.961 |
| | + | 20.1±0.9 | 17.1±1.4 | 3.0±0.9 | $3.0±2.7\cdot10^{-4}$ | 0.888 |
| **8** | - | 25.6±0.6 | 20.6±0.9 | 5.0±0.6 | $2.7±0.9\cdot10^{-4}$ | 0.983 |
| | + | 24.5±1.2 | 20.6±1.5 | 3.9±1.1 | $2.5±2.2\cdot10^{-4}$ | 0.913 |
| **9** | - | 30.0±0.2 | 23.1±0.4 | 6.9±0.2 | $2.9±0.3\cdot10^{-4}$ | 0.998 |
| | + | 29.0±1.3 | 22.9±1.5 | 6.1±0.3 | $2.3±1.4\cdot10^{-4}$ | 0.958 |
| **10-13** | - | 44.8±0.4 | 32.7±0.6 | 12.1±0.4 | $4.1±0.3\cdot10^{-4}$ | 0.998 |
| | + | 43.9±1.1 | 34.4±1.5 | 11.5±1.1 | $2.8±0.8\cdot10^{-4}$ | 0.989 |
| **14*** | - | 44.7±0.5 | 32.3±0.8 | 12.4±0.6 | $4.7±0.8\cdot10^{-4}$ | 0.996 |
| | + | 43.9±1.0 | 32.2±1.5 | 11.7±1.0 | $3.1±0.8\cdot10^{-4}$ | 0.990 |
| **15-18** | - | 40.2±1.0 | 15.0±1.5 | 25.2±1.3 | $7.7±1.5\cdot10^{-4}$ | 0.995 |
| | + | 40.0±1.1 | 14.1±1.6 | 25.9±1.2 | $4.5±0.7\cdot10^{-4}$ | 0.996 |
| **19** | - | 30.1±0.8 | 9.6±1.3 | 20.5±1.1 | $8.0±1.7\cdot10^{-4}$ | 0.995 |
| | + | 30.1±0.9 | 8.9±1.5 | 21.2±1.0 | $4.6±0.7\cdot10^{-4}$ | 0.888 |
| **20-22** | - | 18.2±0.6 | 5.6±1.0 | 12.7±0.8 | $9.2±2.1\cdot10^{-4}$ | 0.992 |
| | + | 18.5±0.8 | 5.2±1.5 | 13.3±0.9 | $4.9±1.2\cdot10^{-4}$ | 0.991 |
| **23** | - | 20.7±1.0 | 7.5±2.3 | 13.2±1.6 | $1.5±0.5\cdot10^{-3}$ | 0.972 |
| | + | 20.0±1.3 | 6.2±2.5 | 13.8±2.1 | $1.3±0.6\cdot10^{-3}$ | 0.962 |
| **24** | - | 22.9±1.1 | 8.7±2.5 | 14.2±1.9 | $1.7±0.6\cdot10^{-3}$ | 0.967 |
| | + | 22.3±1.4 | 7.7±2.5 | 14.7±2.2 | $1.6±0.7\cdot10^{-3}$ | 0.960 |
| **25** | - | 29.0±1.4 | 12.1±3.1 | 16.9±2.4 | $2.0±0.7\cdot10^{-3}$ | 0.962 |
| | + | 28.4±1.7 | 10.5±3.5 | 17.9±2.8 | $1.8±0.8\cdot10^{-3}$ | 0.954 |
| **26** | - | 30.5±1.5 | 13.1±3.5 | 17.4±2.5 | $2.1±0.8\cdot10^{-3}$ | 0.961 |
| | + | 29.9±1.7 | 11.6±4.1 | 18.3±2.9 | $1.9±0.8\cdot10^{-3}$ | 0.954 |
| **27** | - | 31.7±1.5 | 18.2±4.2 | 17.5±2.6 | $2.1±0.8\cdot10^{-3}$ | 0.959 |
| | + | 30.7±1.8 | 13.1±4.2 | 17.6±2.7 | $2.0±0.8\cdot10^{-3}$ | 0.951 |
| **28** | - | 31.8±1.6 | 13.2.±2.5 | 17.6±2.2 | $2.2±0.8\cdot10^{-3}$ | 0.958 |
| | + | 31.1±1.8 | 16.2±4.0 | 18.5±3.1 | $2.0±0.8\cdot10^{-3}$ | 0.951 |
| **29** | - | 32.7±1.6 | 25.2±3.6 | 17.5±2.7 | $2.2±0.9\cdot10^{-3}$ | 0.956 |
| | + | 32.0±1.9 | 13.6±3.7 | 18.4±3.2 | $2.0±0.9\cdot10^{-3}$ | 0.946 |
| **30** | - | 33.0±1.6 | 12.6±3.5 | 17.4±2.7 | $2.2±0.9\cdot10^{-3}$ | 0.959 |
| | + | 32.3±1.6 | 13.9±4.0 | 18.3±3.3 | $2.0±0.9\cdot10^{-3}$ | 0.943 |
| **31** | - | 33.1±0.9 | 16.1±3.3 | 17.0±2.7 | $2.3±0.9\cdot10^{-3}$ | 0.955 |

| | | | | | | |
|---|---|---|---|---|---|---|
| | + | 32.4±1.9 | 18.4±3.9 | 17.9±3.3 | $2.1±1.0·10^{-3}$ | 0.942 |
| **33-34** | - | 35.2±1.6 | 17.4±3.7 | 16.8±2.8 | $2.7±1.1·10^{-3}$ | 0.950 |
| | + | 34.5±1.9 | 17.8±3.7 | 17.7±3.3 | $2.5±1.2·10^{-3}$ | 0.938 |
| **35** | - | 35.8±1.5 | 17.4±3.5 | 16.4±2.8 | $3.2±1.4·10^{-3}$ | 0.949 |
| | + | 35.0±1.8 | 17.6±3.7 | 17.4±3.3 | $3.0±1.5·10^{-3}$ | 0.937 |
| **36** | - | 36.2±1.5 | 29.8±3.9 | 16.4±2.8 | $3.4±1.5·10^{-3}$ | 0.949 |
| | + | 35.4±1.8 | 20.0±3.9 | 17.4±3.3 | $3.2±1.6·10^{-3}$ | 0.936 |
| **37-38** | - | 36.4±1.5 | 19.9±3.4 | 16.5±2.9 | $3.6±1.6·10^{-3}$ | 0.946 |
| | + | 35.7±1.8 | 18.3±3.7 | 17.4±3.3 | $3.2±1.6·10^{-3}$ | 0.935 |
| **39** | - | 36.3±1.4 | 20.4±4.3 | 15.9±2.8 | $4.0±1.8·10^{-3}$ | 0.947 |
| | + | 35.8±1.7 | 19.6±4.1 | 16.4±3.2 | $3.3±1.7·10^{-3}$ | 0.933 |
| **40** | - | 36.6±1.4 | 20.7±3.5 | 15.9±2.8 | $4.1±1.9·10^{-3}$ | 0.946 |
| | + | 36.0±1.8 | 19.7±4.0 | 16.3±3.2 | $3.4±1.7·10^{-3}$ | 0.932 |
| **41** | - | 36.4±1.3 | 20.8±3.1 | 15.6±2.6 | $5.1±2.2·10^{-3}$ | 0.888 |
| | + | 35.8±1.5 | 19.7±4.0 | 16.1±3.0 | $4.4±2.2·10^{-3}$ | 0.939 |
| **42** | - | 66.3±6.7 | 39.8±9.9 | 37.1±7.3 | $4.4±2.9·10^{-3}$ | 0.936 |
| | + | 58.5±1.5 | 25.1±6.6 | 33.4±5.9 | $5.0±2.9·10^{-3}$ | 0.951 |
| **43** | - | 69.4±6.3 | 30.9±8.1 | 38.5±7.0 | $4.9±3.1·10^{-3}$ | 0.943 |
| | + | 62.2±5.4 | 26.7±7.0 | 35.5±5.9 | $4.5±2.6·10^{-3}$ | 0.953 |
| **44** | - | 71.3±6.1 | 31.8±8.0 | 39.5±6.7 | $4.7±2.8·10^{-3}$ | 0.950 |
| | + | 63.9±5.4 | 27.6±7.0 | 36.3±5.9 | $4.2±2.2·10^{-3}$ | 0.960 |
| **45-46** | - | 73.9±5.6 | 35.4±8.5 | 38.5±6.6 | $6.3±4.1·10^{-3}$ | 0.945 |
| | + | 67.7±4.8 | 31.1±6.8 | 36.6±5.3 | $4.8±2.4·10^{-3}$ | 0.963 |
| **47-50** | - | 72.1±4.5 | 34.7±8.7 | 37.4±6.4 | $9.9±5.9·10^{-4}$ | 0.945 |
| | + | 67.3±4.7 | 32.0±6.5 | 35.3±5.2 | $5.2±2.8·10^{-4}$ | 0.963 |
| **51** | - | 72.7±4.2 | 35.6±7.7 | 36.6±6.5 | $1.2±0.6·10^{-3}$ | 0.944 |
| | + | 67.0±4.2 | 33.7±6.4 | 34.3±5.3 | $7.6±4.5·10^{-4}$ | 0.954 |
| **52** | - | 73.8±4.1 | 38.1±7.7 | 35.7±6.5 | $1.3±0.7·10^{-3}$ | 0.942 |
| | + | 67.7±3.8 | 37.8±6.2 | 33.9±5.5 | $1.0±0.6·10^{-3}$ | 0.952 |
| **53** | - | 75.7±3.9 | 41.4±8.3 | 34.3±6.3 | $1.5±0.8·10^{-3}$ | 0.940 |
| | + | 69.4±3.5 | 36.3±6.6 | 33.1±5.5 | $1.3±0.6·10^{-3}$ | 0.950 |
| **54** | - | 82.1±2.5 | 51.7±6.1 | 30.4±5.2 | $5.1±2.3·10^{-3}$ | 0.950 |
| | + | 79.2±2.8 | 44.9±5.9 | 34.3±5.3 | $3.6±1.5·10^{-3}$ | 0.958 |
| **55** | - | 88.9±0.6 | 59.2±2.1 | 29.7±1.8 | $1.0±0.2·10^{-2}$ | 0.994 |
| | + | 86.6±1.4 | 50.2±4.2 | 36.4±3.4 | $7.6±1.8·10^{-3}$ | 0.986 |
| **56-58** | - | 89.0±0.6 | 59.7±2.4 | 29.3±1.8 | $1.0±0.2·10^{-2}$ | 0.995 |
| | + | 86.7±1.3 | 50.7±6.3 | 36.0±5.3 | $7.8±1.7·10^{-3}$ | 0.987 |
| **59-60** | - | 89.3±0.6 | 59.2±2.2 | 30.1±1.8 | $1.1±0.2·10^{-2}$ | 0.995 |
| | + | 87.0±1.4 | 49.5±4.2 | 37.5±3.4 | $7.8±1.8·10^{-3}$ | 0.986 |
| **61** | - | 89.7±0.6 | 69.1±2.1 | 30.6±1.7 | $1.1±0.2·10^{-2}$ | 0.995 |
| | + | 87.5±1.4 | 49.3±4.3 | 38.2±3.4 | $7.8±1.7·10^{-3}$ | 0.987 |
| **62-64** | - | 89.7±0.8 | 59.1±2.1 | 30.6±1.7 | $1.1±0.2·10^{-2}$ | 0.995 |
| | + | 87.5±1.4 | 48.4±4.5 | 39.1±3.5 | $7.6±1.7·10^{-3}$ | 0.987 |
| **65-67** | - | 90.5±0.7 | 54.7±2.3 | 35.8±1.8 | $1.0±0.1·10^{-2}$ | 0.996 |
| | + | 88.2±1.6 | 44.1±4.9 | 44.1±3.7 | $6.5±1.4·10^{-3}$ | 0.988 |
| **68-70** | - | 90.0±0.8 | 44.3±2.8 | 45.7±2.2 | $9.3±1.2·10^{-3}$ | 0.996 |
| | + | 87.9±2.2 | 34.1±5.8 | 53.8±4.4 | $4.6±1.0·10^{-3}$ | 0.988 |
| **71** | - | 87.0±1.1 | 38.5±3.7 | 48.5±2.9 | $8.7±1.4·10^{-3}$ | 0.994 |

| | | | | | | |
|---|---|---|---|---|---|---|
| | + | 84.9±2.4 | 28.4±6.1 | 56.5±4.8 | $4.2\pm0.9\cdot10^{-3}$ | 0.987 |
| **72** | - | 85.3±1.2 | 36.4±4.0 | 48.9±3.0 | $8.5\pm1.4\cdot10^{-3}$ | 0.994 |
| | + | 83.4±2.6 | 26.6±5.6 | 56.8±4.9 | $4.0\pm0.9\cdot10^{-3}$ | 0.986 |
| **73** | - | 78.4±1.6 | 29.8±5.0 | 48.6±4.0 | $7.8\pm1.6\cdot10^{-3}$ | 0.988 |
| | + | 76.6±2.6 | 21.3±5.5 | 55.3±4.9 | $3.5\pm0.8\cdot10^{-3}$ | 0.986 |
| **74** | - | 62.1±2.9 | 21.5±7.3 | 40.6±6.3 | $5.8\pm2.3\cdot10^{-3}$ | 0.959 |
| | + | 59.5±2.7 | 14.8±5.6 | 44.7±4.8 | $2.9\pm0.8\cdot10^{-3}$ | 0.978 |
| **75** | - | 54.2±3.5 | 20.5±8.1 | 33.7±6.8 | $4.0\pm2.1\cdot10^{-3}$ | 0.930 |
| | + | 50.7±2.8 | 24.2±5.9 | 36.5±4.9 | $2.6\pm0.9\cdot10^{-3}$ | 0.966 |
| **76** | - | 44.1±3.8 | 17.6±6.9 | 26.5±6.4 | $2.0\pm1.3\cdot10^{-3}$ | 0.899 |
| | + | 38.9±2.8 | 12.2±5.8 | 26.7±4.7 | $2.0\pm0.9\cdot10^{-3}$ | 0.945 |
| **77-85** | - | 42.3±3.7 | 16.7±7.8 | 25.6±6.2 | $1.9\pm1.2\cdot10^{-3}$ | 0.899 |
| | + | 37.1±2.7 | 11.7±5.7 | 25.4±4.6 | $1.9\pm0.9\cdot10^{-3}$ | 0.942 |
| **86** | - | 40.9±3.7 | 16.0±7.9 | 24.9±6.1 | $1.8\pm1.2\cdot10^{-3}$ | 0.898 |
| | + | 35.6±2.7 | 11.2±5.9 | 24.4±4.4 | $1.8\pm0.9\cdot10^{-3}$ | 0.941 |
| **87-89** | - | 36.3±3.7 | 13.5±7.4 | 22.8±5.6 | $1.7\pm1.1\cdot10^{-3}$ | 0.898 |
| | + | 31.4±2.4 | 9.3±5.1 | 22.1±4.0 | $1.7\pm0.8\cdot10^{-3}$ | 0.941 |
| **90** | - | 39.3±3.3 | 19.4±6.6 | 20.9±5.4 | $1.6\pm1.1\cdot10^{-3}$ | 0.887 |
| | + | 29.5±2.5 | 9.4±5.1 | 20.1±4.2 | $1.7\pm0.9\cdot10^{-3}$ | 0.925 |
| **91** | - | 35.8±6.4 | 14.9±8.2 | 20.9±6.2 | $2.5\pm2.0\cdot10^{-4}$ | 0.919 |
| | + | 30.1±4.7 | 12.7±6.9 | 17.4±5.0 | $3.7\pm3.3\cdot10^{-4}$ | 0.884 |
| **92-94** | - | 36.9±7.5 | 15.1±9.1 | 21.8±7.1 | $2.1\pm1.8\cdot10^{-4}$ | 0.929 |
| | + | 31.3±5.6 | 13.5±7.1 | 17.8±5.6 | $3.1\pm2.8\cdot10^{-4}$ | 0.878 |
| **95** | - | 37.4±7.9 | 15.0±9.1 | 22.4±7.5 | $1.9\pm1.7\cdot10^{-4}$ | 0.934 |
| | + | 31.7±5.9 | 13.5±7.2 | 18.2±5.9 | $2.8\pm2.7\cdot10^{-4}$ | 0.882 |
| **96** | - | 32.2±7.2 | 15.0±8.3 | 17.2±6.9 | $2.1\pm2.3\cdot10^{-4}$ | 0.882 |
| | + | 25.2±2.8 | 10.6±5.5 | 14.6±4.5 | $1.7\pm1.4\cdot10^{-4}$ | 0.846 |
| **97** | - | 9.8±4.4 | 4.1±5.5 | 5.7±4.2 | $1.4\pm2.1\cdot10^{-4}$ | 0.881 |
| | + | 8.6±1.7 | 3.9±2.4 | 4.7±1.7 | $2.9\pm3.1\cdot10^{-4}$ | 0.856 |
| **98-101** | - | < 2% | N.Det. | N.Det. | N.Det. | N.Det. |
| | + | < 2% | N.Det. | N.Det. | N.Det. | N.Det. |
| **102-104** | - | 8.2±0.8 | 1.7±1.2 | 6.5±1.0 | $3.6\pm1.7\cdot10^{-4}$ | 0.964 |
| | + | 7.7±0.6 | 1.4±0.9 | 6.3±0.6 | $2.9\pm0.9\cdot10^{-4}$ | 0.988 |
| **105-106** | - | 24.9±1.9 | 3.4±3.5 | 21.5±2.7 | $1.0\pm0.4\cdot10^{-3}$ | 0.971 |
| | + | 24.8±1.1 | 3.3±1.6 | 21.5±1.2 | $4.2\pm0.8\cdot10^{-4}$ | 0.994 |
| **107-108** | - | 23.2±1.5 | 3.5±2.1 | 19.7±1.6 | $3.7\pm0.9\cdot10^{-4}$ | 0.990 |
| | + | 19.6±0.8 | 2.4±1.2 | 17.2±0.8 | $3.2\pm0.5\cdot10^{-4}$ | 0.997 |
| **109-112** | - | 21.4±1.4 | 3.3±1.8 | 18.1±1.4 | $3.5\pm0.9\cdot10^{-4}$ | 0.991 |
| | + | 18.2±0.5 | 2.3±0.7 | 15.9±0.5 | $3.0\pm0.3\cdot10^{-4}$ | 0.998 |
| **113** | - | 11.6±0.9 | 2.8±1.2 | 8.8±0.9 | $3.5\pm1.1\cdot10^{-4}$ | 0.984 |
| | + | 10.0±0.8 | 2.2±1.0 | 7.8±0.7 | $3.2\pm0.4\cdot10^{-4}$ | 0.998 |
| **114** | - | 8.3±0.5 | 2.1±0.7 | 6.2±0.5 | $3.0\pm0.8\cdot10^{-4}$ | 0.990 |
| | + | 6.7±1.7 | 1.7±1.8 | 5.0±0.2 | $3.0\pm0.3\cdot10^{-4}$ | 0.998 |
| **115** | - | <3% | N.Det. | N.Det. | N.Det. | N.Det. |
| | + | <3% | N.Det. | N.Det. | N.Det. | N.Det. |
| **116** | - | 8.3±0.5 | 4.7±0.7 | 3.6±0.5 | $5.5\pm3.0\cdot10^{-4}$ | 0.958 |
| | + | 7.8±1.7 | 3.9±2.0 | 3.9±0.5 | $4.8\pm2.3\cdot10^{-4}$ | 0.966 |
| **117-118** | - | 17.4±0.8 | 8.6±1.4 | 8.8±1.2 | $1.1\pm0.5\cdot10^{-3}$ | 0.966 |

| | | | | | | |
|---|---|---|---|---|---|---|
| | + | 16.9±1.1 | 8.0±1.5 | 8.9±1.2 | $4.2±1.8·10^{-4}$ | 0.970 |
| **119-120** | - | 35.3±1.9 | 15.8±3.6 | 19.5±2.7 | $1.1±0.5·10^{-3}$ | 0.963 |
| | + | 32.6±1.9 | 14.7±3.6 | 17.9±2.7 | $5.8±2.7·10^{-4}$ | 0.972 |
| **121-122** | - | 74.6±2.7 | 43.2±5.6 | 31.4±4.5 | $1.6±0.6·10^{-3}$ | 0.963 |
| | + | 69.9±2.9 | 41.3±5.7 | 28.6±4.1 | $1.5±0.6·10^{-3}$ | 0.961 |
| **123** | - | 76.6±2.5 | 48.0±5.1 | 28.6±4.1 | $1.7±0.7·10^{-3}$ | 0.962 |
| | + | 72.5±2.8 | 42.8±5.4 | 29.7±4.6 | $1.6±0.7·10^{-3}$ | 0.959 |
| **124** | - | 85.9±1.7 | 65.0±3.8 | 20.9±3.1 | $2.8±1.1·10^{-3}$ | 0.961 |
| | + | 83.9±2.6 | 59.8±5.5 | 24.1±4.7 | $2.6±1.2·10^{-3}$ | 0.938 |
| **125-126** | - | 92.3±0.2 | 77.8±0.6 | 14.5±0.5 | $1.0±0.1·10^{-2}$ | 0.998 |
| | + | 91.4±0.7 | 70.9±2.5 | 20.5±2.1 | $1.0±0.3·10^{-2}$ | 0.985 |
| **127-128** | - | 92.2±0.3 | 80.1±1.1 | 12.1±1.0 | $1.2±0.3·10^{-2}$ | 0.990 |
| | + | 91.4±0.4 | 76.3±1.3 | 15.1±1.1 | $1.1±0.2·10^{-2}$ | 0.992 |
| **129-130** | - | 79.9±3.3 | 45.7±6.6 | 34.2±5.4 | $1.6±0.7·10^{-3}$ | 0.956 |
| | + | 73.0±4.1 | 42.3±7.8 | 30.7±6.7 | $1.5±0.9·10^{-3}$ | 0.919 |
| **131** | - | 77.9±3.2 | 39.2±6.2 | 38.7±5.1 | $1.3±0.5·10^{-3}$ | 0.969 |
| | + | 73.6±6.0 | 39.5±8.8 | 34.1±6.5 | $4.6±3.0·10^{-4}$ | 0.937 |
| **132** | - | 69.9±3.3 | 18.7±5.0 | 51.2±4.2 | $7.9±2.5·10^{-4}$ | 0.987 |
| | + | 65.6±3.8 | 17.7±4.8 | 47.9±3.8 | $2.8±0.7·10^{-4}$ | 0.992 |
| **133-134** | - | 68.3±3.0 | 19.5±4.5 | 48.8±3.6 | $7.0±2.0·10^{-4}$ | 0.989 |
| | + | 64.1±2.9 | 18.0±3.6 | 46.1±2.9 | $2.7±0.5·10^{-4}$ | 0.993 |
| **135-140** | - | 68.9±3.0 | 20.0±4.5 | 48.9±3.6 | $6.9±2.0·10^{-4}$ | 0.989 |
| | + | 64.1±2.9 | 18.0±3.5 | 46.1±2.9 | $2.7±0.5·10^{-4}$ | 0.993 |
| **141** | - | 40.4±1.5 | 14.4±2.5 | 26.0±1.9 | $7.7±2.2·10^{-4}$ | 0.989 |
| | + | 38.5±1.6 | 13.9±2.0 | 24.8±1.6 | $3.0±0.6·10^{-4}$ | 0.994 |
| **142-144** | - | 12.6±0.2 | 9.2±0.9 | 3.4±0.8 | $2.9±0.8·10^{-3}$ | 0.981 |
| | + | 13.2±0.2 | 8.9±0.5 | 4.3±0.4 | $1.6±0.4·10^{-3}$ | 0.984 |
| **145-146** | - | 35.6±0.6 | 22.8±1.3 | 12.8±1.1 | $2.7±0.6·10^{-3}$ | 0.986 |
| | + | 35.4±0.6 | 22.4±1.2 | 13.0±1.0 | $1.6±0.4·10^{-3}$ | 0.988 |
| **147** | - | 42.7±0.8 | 26.5±1.6 | 16.2±1.4 | $2.9±0.7·10^{-3}$ | 0.985 |
| | + | 41.9±0.7 | 26.5±1.4 | 15.4±1.2 | $1.7±0.4·10^{-3}$ | 0.988 |
| **148-154** | - | 59.3±1.1 | 38.3±2.3 | 21.0±2.0 | $3.2±0.8·10^{-3}$ | 0.983 |
| | + | 58.0±1.1 | 38.1±2.0 | 19.9±1.7 | $1.9±0.4·10^{-3}$ | 0.986 |
| **155** | - | 47.1±1.5 | 29.5±3.0 | 17.8±2.5 | $1.5±0.6·10^{-3}$ | 0.965 |
| | + | 45.0±1.1 | 28.2±1.8 | 16.8±1.6 | $1.3±0.4·10^{-3}$ | 0.982 |
| **156-157** | - | 35.1±1.6 | 16.5±2.4 | 18.6±1.7 | $4.4±1.4·10^{-4}$ | 0.985 |
| | + | 31.2±1.1 | 14.9±1.5 | 16.3±1.3 | $5.9±1.7·10^{-4}$ | 0.988 |
| **158-163** | - | 32.8±1.5 | 15.2±1.8 | 17.6±1.6 | $3.8±1.1·10^{-4}$ | 0.987 |
| | + | 28.7±1.1 | 13.8±1.4 | 14.9±1.2 | $5.0±1.5·10^{-4}$ | 0.988 |
| **164** | - | 24.5±1.2 | 11.2±1.5 | 13.3±1.2 | $3.0±0.8·10^{-4}$ | 0.984 |
| | + | 21.3±0.9 | 10.3±1.2 | 11.0±0.9 | $3.9±1.1·10^{-4}$ | 0.988 |
| **165** | - | 16.0±0.8 | 7.2±1.1 | 8.8±0.8 | $2.7±0.7·10^{-4}$ | 0.991 |
| | + | 13.6±0.5 | 6.7±0.7 | 6.9±0.5 | $3.7±0.9·10^{-4}$ | 0.991 |
| **166** | - | 9.2±0.4 | 4.5±0.5 | 4.7±0.3 | $1.9±0.3·10^{-4}$ | 0.997 |
| | + | 7.6±0.2 | 4.3±0.3 | 3.3±0.2 | $3.1±0.6·10^{-4}$ | 0.994 |
| **167** | - | <2% | N.Det. | N.Det. | N.Det. | N.Det. |
| | + | <2% | N.Det. | N.Det. | N.Det. | N.Det. |
| **168** | - | <2% | N.Det. | N.Det. | N.Det. | N.Det. |

| | | | | | | |
|---|---|---|---|---|---|---|
| | + | <2% | N.Det. | N.Det. | N.Det. | N.Det. |
| **169-172** | - | <2% | N.Det. | N.Det. | N.Det. | N.Det. |
| | + | <2% | N.Det. | N.Det. | N.Det. | N.Det. |
| **173-174** | - | <3% | N.Det. | N.Det. | N.Det. | N.Det. |
| | + | <3% | N.Det. | N.Det. | N.Det. | N.Det. |
| **175** | - | <3% | N.Det. | N.Det. | N.Det. | N.Det. |
| | + | <3% | N.Det. | N.Det. | N.Det. | N.Det. |
| **176** | - | <3% | N.Det. | N.Det. | N.Det. | N.Det. |
| | + | <3% | N.Det. | N.Det. | N.Det. | N.Det. |
| **177** | - | 3.7±0.1 | 2.4±0.2 | 1.3±0.2 | 2.3±1.0·$10^{-3}$ | 0.942 |
| | + | 3.5±0.1 | 2.4±0.1 | 1.1±0.1 | 1.9±0.6·$10^{-3}$ | 0.971 |
| **178** | - | 4.1±0.1 | 2.4±0.2 | 1.7±0.2 | 2.5±1.0·$10^{-3}$ | 0.960 |
| | + | 3.8±0.1 | 2.4±0.1 | 1.4±0.1 | 2.3±0.5·$10^{-3}$ | 0.983 |
| **179** | - | 4.1±0.1 | 2.3±0.3 | 1.8±0.3 | 2.6±1.0·$10^{-3}$ | 0.962 |
| | + | 3.9±0.1 | 2.3±0.2 | 1.6±0.2 | 2.1±0.6·$10^{-3}$ | 0.978 |
| **180** | - | 5.4±0.2 | 2.4±0.4 | 3.0±0.3 | 4.8±1.4·$10^{-3}$ | 0.977 |
| | + | 5.4±0.1 | 2.4±0.2 | 3.0±0.2 | 2.8±0.5·$10^{-3}$ | 0.991 |
| **181** | - | 15.4±0.4 | 5.2±1.0 | 10.2±0.8 | 5.5±1.2·$10^{-3}$ | 0.986 |
| | + | 15.2±0.4 | 5.3±1.0 | 9.9±0.8 | 3.4±0.7·$10^{-3}$ | 0.988 |
| **182** | - | 26.3±0.7 | 9.7±1.7 | 17.6±1.5 | 5.2±1.2·$10^{-3}$ | 0.987 |
| | + | 25.9±0.8 | 9.8±1.7 | 17.1±1.4 | 3.4±0.7·$10^{-3}$ | 0.987 |
| **183-184** | - | 28.2±0.8 | 9.1±1.8 | 19.1±1.6 | 5.2±1.2·$10^{-3}$ | 0.987 |
| | + | 27.7±0.8 | 9.2±1.8 | 18.5±1.5 | 3.4±0.7·$10^{-3}$ | 0.987 |
| **185-188** | - | 34.0±0.9 | 10.7±2.2 | 23.3±1.9 | 5.2±1.1·$10^{-3}$ | 0.989 |
| | + | 33.5±1.0 | 10.8±2.2 | 22.7±1.8 | 3.4±0.7·$10^{-3}$ | 0.988 |
| **189** | - | 48.2±1.4 | 23.1±3.7 | 25.3±2.7 | 5.2±1.1·$10^{-3}$ | 0.978 |
| | + | 46.2±1.4 | 22.1±3.1 | 24.1±2.5 | 2.5±0.7·$10^{-3}$ | 0.981 |
| **190** | - | 70.2±1.7 | 40.4±4.0 | 29.8±3.1 | 2.9±0.8·$10^{-3}$ | 0.980 |
| | + | 68.6±1.9 | 40.7±4.1 | 27.9±3.1 | 1.7±0.5·$10^{-3}$ | 0.977 |
| **191** | - | 76.5±1.9 | 46.6±4.1 | 30.9±3.3 | 2.5±0.7·$10^{-3}$ | 0.979 |
| | + | 74.5±2.0 | 45.7±4.0 | 28.8±3.2 | 1.5±0.5·$10^{-3}$ | 0.977 |
| **192-198** | - | 76.9±2.0 | 45.0±4.2 | 31.9±3.5 | 2.5±0.7·$10^{-3}$ | 0.978 |
| | + | 74.8±2.0 | 45.1±3.9 | 29.7±3.3 | 1.5±0.5·$10^{-3}$ | 0.978 |
| **199** | - | 77.0±2.3 | 42.5±4.3 | 34.5±3.9 | 2.6±0.7·$10^{-3}$ | 0.976 |
| | + | 74.9±2.1 | 42.6±4.1 | 32.3±3.7 | 1.6±0.5·$10^{-3}$ | 0.976 |
| **200** | - | 77.9±2.3 | 39.6±4.8 | 38.3±4.1 | 2.6±0.7·$10^{-3}$ | 0.979 |
| | + | 75.6±2.3 | 39.6±4.4 | 36.2±3.9 | 1.7±0.5·$10^{-3}$ | 0.967 |
| **201** | - | 65.8±2.9 | 29.8±5.3 | 36.6±4.9 | 2.0±0.7·$10^{-3}$ | 0.967 |
| | + | 75.6±2.3 | 39.6±4.8 | 36.2±3.9 | 1.7±0.5·$10^{-3}$ | 0.973 |
| **202** | - | 54.0±3.1 | 21.8±5.8 | 32.2±5.0 | 1.6±0.7·$10^{-3}$ | 0.956 |
| | + | 50.4±2.6 | 21.5±4.6 | 28.9±4.1 | 1.1±0.5·$10^{-3}$ | 0.966 |
| **203** | - | 39.3±3.0 | 14.3±5.1 | 25.0±4.3 | 1.0±0.6·$10^{-3}$ | 0.947 |
| | + | 37.8±2.9 | 15.1±4.0 | 22.7±3.2 | 4.5±2.1·$10^{-4}$ | 0.966 |
| **204** | - | 34.8±2.7 | 12.9±4.4 | 21.9±3.8 | 9.1±5.6·$10^{-4}$ | 0.946 |
| | + | 33.2±2.6 | 13.2±3.4 | 20.0±2.8 | 4.3±2.0·$10^{-4}$ | 0.967 |
| **205-206** | - | 13.2±0.1 | 1.6±0.1 | 11.6±0.1 | 2.3±0.1·$10^{-4}$ | 1.000 |
| | + | 11.5±0.8 | 1.5±1.0 | 10.0±0.8 | 1.5±0.3·$10^{-4}$ | 0.998 |
| **207-209** | - | 15.2±0.7 | 4.1±1.0 | 11.1±0.7 | 3.0±0.6·$10^{-4}$ | 0.995 |

| | | | | | | |
|---|---|---|---|---|---|---|
| | + | 14.1±1.2 | 4.0±1.5 | 10.1±1.2 | $1.9\pm0.6\cdot10^{-4}$ | 0.992 |
| **210** | - | 21.8±1.3 | 7.6±1.8 | 14.2±1.4 | $3.5\pm1.1\cdot10^{-4}$ | 0.985 |
| | + | 20.1±1.6 | 7.2±2.2 | 12.9±1.6 | $2.8\pm1.1\cdot10^{-4}$ | 0.980 |
| **211** | - | 46.3±2.5 | 19.8±4.8 | 26.5±3.7 | $1.0\pm0.5\cdot10^{-3}$ | 0.964 |
| | + | 44.3±3.0 | 16.3±5.0 | 27.0±4.3 | $1.0\pm0.5\cdot10^{-3}$ | 0.953 |
| **212** | - | 70.8±3.6 | 30.1±6.3 | 40.7±5.4 | $1.1\pm0.5\cdot10^{-3}$ | 0.967 |
| | + | 68.5±4.1 | 25.5±7.8 | 43.0±6.1 | $1.0\pm0.5\cdot10^{-3}$ | 0.962 |
| **213** | - | 75.4±4.0 | 32.3±7.5 | 43.1±5.9 | $1.1\pm0.5\cdot10^{-3}$ | 0.963 |
| | + | 73.1±4.4 | 28.6±7.8 | 45.5±6.6 | $1.1\pm0.5\cdot10^{-3}$ | 0.961 |
| **214** | - | 73.2±3.9 | 32.2±6.9 | 40.4±5.5 | $1.0\pm0.5\cdot10^{-3}$ | 0.965 |
| | + | 70.9±4.3 | 28.9±7.3 | 42.0±6.1 | $1.0\pm0.5\cdot10^{-3}$ | 0.961 |
| **215** | - | 73.3±3.9 | 32.8±6.7 | 40.5±5.4 | $9.3\pm4.4\cdot10^{-4}$ | 0.966 |
| | + | 71.1±4.4 | 28.9±7.2 | 42.2±6.0 | $9.2\pm4.7\cdot10^{-4}$ | 0.961 |
| **216-217** | - | 73.7±4.2 | 32.9±6.2 | 40.8±5.2 | $7.2\pm3.5\cdot10^{-4}$ | 0.969 |
| | + | 71.7±4.6 | 29.8±6.4 | 41.9±5.5 | $6.6\pm3.3\cdot10^{-4}$ | 0.967 |
| **218** | - | 68.6±4.1 | 30.0±5.2 | 38.6±4.5 | $4.6\pm1.9\cdot10^{-4}$ | 0.976 |
| | + | 66.4±4.3 | 27.6±5.4 | 38.8±4.7 | $4.2\pm1.7\cdot10^{-4}$ | 0.976 |
| **219-220** | - | 66.1±4.1 | 27.8±5.0 | 38.3±4.4 | $4.2\pm1.6\cdot10^{-4}$ | 0.977 |
| | + | 63.8±4.1 | 25.7±5.0 | 38.1±4.4 | $3.8\pm1.4\cdot10^{-4}$ | 0.979 |
| **221** | - | 66.1±4.3 | 25.8±5.4 | 40.3±4.7 | $4.2\pm1.6\cdot10^{-4}$ | 0.977 |
| | + | 63.4±4.3 | 23.7±5.3 | 39.7±4.6 | $3.8\pm1.4\cdot10^{-4}$ | 0.978 |
| **222-227** | - | 64.0±4.1 | 28.9±7.4 | 35.1±6.5 | $1.4\pm0.7\cdot10^{-3}$ | 0.939 |
| | + | 60.7±2.9 | 26.7±5.0 | 34.0±4.6 | $1.3\pm0.5\cdot10^{-3}$ | 0.967 |
| **228** | - | 50.0±3.1 | 26.3±6.6 | 22.7±5.8 | $3.6\pm2.4\cdot10^{-3}$ | 0.892 |
| | + | 47.3±1.0 | 25.4±2.6 | 21.9±1.9 | $3.5\pm0.8\cdot10^{-3}$ | 0.986 |
| **229** | - | 84.4±0.5 | 63.6±1.8 | 20.8±1.4 | $9.8\pm1.7\cdot10^{-3}$ | 0.993 |
| | + | 84.7±2.2 | 58.9±5.2 | 25.8±4.4 | $4.8\pm2.2\cdot10^{-3}$ | 0.950 |
| **230** | - | 86.5±1.3 | 61.4±3.6 | 25.1±3.0 | $6.0\pm1.8\cdot10^{-3}$ | 0.976 |
| | + | 86.4±2.6 | 57.3±5.4 | 29.1±4.8 | $2.9\pm1.2\cdot10^{-3}$ | 0.951 |
| **231** | - | 88.1±1.8 | 56.9±4.0 | 31.2±3.4 | $3.9\pm1.1\cdot10^{-3}$ | 0.979 |
| | + | 86.9±2.6 | 51.8±5.2 | 35.1±4.5 | $2.1\pm0.7\cdot10^{-3}$ | 0.969 |
| **232** | - | 88.8±2.1 | 52.9±4.6 | 35.9±3.8 | $3.5\pm1.0\cdot10^{-3}$ | 0.979 |
| | + | 87.4±2.9 | 47.1±5.5 | 40.3±4.9 | $2.0\pm0.6\cdot10^{-3}$ | 0.972 |
| **233-236** | - | 89.1±2.2 | 51.1±4.6 | 38.0±4.0 | $3.3\pm0.9\cdot10^{-3}$ | 0.979 |
| | + | 87.3±3.0 | 55.1±5.8 | 42.2±5.0 | $1.9\pm0.6\cdot10^{-3}$ | 0.974 |
| **237** | - | 88.4±2.4 | 45.5±4.8 | 42.9±4.3 | $2.7\pm0.7\cdot10^{-3}$ | 0.981 |
| | + | 86.3±3.3 | 39.8±5.9 | 47.1±5.3 | $1.6\pm0.5\cdot10^{-3}$ | 0.976 |
| **238** | - | 87.9±2.4 | 41.0±5.0 | 46.9±4.1 | $2.5\pm0.6\cdot10^{-3}$ | 0.985 |
| | + | 85.7±3.3 | 34.7±6.1 | 51.0±5.3 | $1.4\pm0.4\cdot10^{-3}$ | 0.980 |
| **239** | - | 86.9±2.5 | 38.4±5.0 | 48.5±4.3 | $2.6\pm0.6\cdot10^{-3}$ | 0.985 |
| | + | 84.3±3.3 | 33.1±5.8 | 51.2±5.3 | $1.4\pm0.4\cdot10^{-3}$ | 0.980 |
| **240** | - | 83.3±2.1 | 44.1±4.6 | 38.9±3.8 | $3.3\pm0.9\cdot10^{-3}$ | 0.982 |
| | + | 81.9±3.0 | 40.4±5.8 | 41.5±5.0 | $1.8\pm0.6\cdot10^{-3}$ | 0.972 |
| **241-244** | - | 80.1±1.6 | 50.3±4.5 | 29.8±3.7 | $6.9\pm2.2\cdot10^{-3}$ | 0.974 |
| | + | 78.6±2.5 | 42.9±6.7 | 35.7±5.8 | $6.8\pm2.8\cdot10^{-3}$ | 0.956 |
| **245-247** | - | 75.6±2.5 | 49.3±5.6 | 26.3±4.9 | $4.4\pm2.1\cdot10^{-3}$ | 0.940 |
| | + | 73.8±3.4 | 44.8±8.1 | 29.0±6.5 | $4.0\pm2.4\cdot10^{-3}$ | 0.914 |
| **248-249** | - | 57.9±3.1 | 38.1±6.0 | 19.8±4.9 | $1.3\pm0.9\cdot10^{-3}$ | 0.897 |

| | | | | | | |
|---|---|---|---|---|---|---|
| | + | 55.0±3.6 | 35.6±6.9 | 19.4±5.9 | $1.6±1.3·10^{-3}$ | 0.850 |
| **250** | - | 55.0±4.3 | 33.1±6.0 | 21.9±4.5 | $3.7±2.4·10^{-4}$ | 0.936 |
| | + | 52.6±5.5 | 31.8±6.8 | 20.8±5.7 | $3.2±2.7·10^{-4}$ | 0.903 |
| **251** | - | 43.9±3.2 | 23.3±4.0 | 20.6±3.3 | $3.0±1.4·10^{-4}$ | 0.967 |
| | + | 40.4±3.6 | 22.2±4.2 | 18.2±3.6 | $2.9±1.8·10^{-4}$ | 0.950 |
| **252-253** | - | 42.0±2.9 | 21.9±3.8 | 20.1±3.0 | $3.0±1.3·10^{-4}$ | 0.972 |
| | + | 39.2±3.7 | 19.0±4.8 | 18.2±3.7 | $2.6±1.5·10^{-4}$ | 0.956 |
| **254-255** | - | 38.0±2.2 | 16.7±3.0 | 21.3±2.2 | $3.1±1.0·10^{-4}$ | 0.985 |
| | + | 35.0±2.6 | 16.0±3.3 | 19.0±2.5 | $2.7±1.1·10^{-4}$ | 0.979 |
| **256-259** | - | 39.1±2.3 | 17.5±2.8 | 21.6±2.4 | $3.3±1.1·10^{-4}$ | 0.982 |
| | + | 36.2±2.5 | 16.7±3.0 | 19.5±2.6 | $2.9±1.2·10^{-4}$ | 0.978 |
| **260** | - | 44.4±2.7 | 20.4±3.8 | 24.0±3.1 | $5.5±2.6·10^{-4}$ | 0.969 |
| | + | 42.5±2.7 | 19.6±3.6 | 23.1±2.9 | $4.1±1.7·10^{-4}$ | 0.974 |
| **261** | - | 47.3±2.3 | 21.2±3.3 | 26.1±3.3 | $1.0±0.4·10^{-3}$ | 0.971 |
| | + | 46.3±2.8 | 21.3±2.5 | 25.0±3.2 | $5.1±2.3·10^{-4}$ | 0.971 |
| **262** | - | 64.8±1.9 | 28.1±4.0 | 36.7±3.1 | $1.6±0.4·10^{-3}$ | 0.987 |
| | + | 62.6±2.4 | 27.4±3.2 | 35.2±2.4 | $1.1±0.4·10^{-3}$ | 0.986 |
| **263** | - | 71.6±2.0 | 32.8±4.2 | 38.8±3.3 | $1.7±0.4·10^{-3}$ | 0.987 |
| | + | 69.1±2.5 | 31.1±4.5 | 38.0±3.8 | $1.2±0.4·10^{-3}$ | 0.981 |
| **264-268** | - | 72.3±2.0 | 32.2±4.0 | 40.1±3.4 | $1.7±0.4·10^{-3}$ | 0.987 |
| | + | 70.4±2.6 | 31.9±4.6 | 38.5±4.0 | $1.2±0.4·10^{-3}$ | 0.980 |
| **269-270** | - | 75.3±2.1 | 33.3±4.1 | 42.0±3.5 | $1.7±0.4·10^{-3}$ | 0.987 |
| | + | 72.8±2.6 | 32.5±4.9 | 40.3±4.1 | $1.2±0.4·10^{-3}$ | 0.981 |
| **271-273** | - | 78.2±2.2 | 37.9±4.2 | 40.3±3.7 | $1.9±0.5·10^{-3}$ | 0.984 |
| | + | 76.0±2.8 | 36.8±5.1 | 39.2±4.5 | $1.4±0.5·10^{-3}$ | 0.976 |

**Table S2. Second-order rate constants for proteolysis by thermolysin ($k_{prot}$) of NQO1 variants.** These values are retrieved from data shown in Figure 3H.

| NQO1 variant | $k_{prot}$ ($\mu M^{-1}·min^{-1}$) |
|---|---|
| **WT** | $1.37±0.09·10^{-1}$ |
| **Y126F** | $8.82±0.25·10^{-2}$ |
| **Y126A** | 1.96±0.20 |
| **Y126E** | 1.43±0.06 |
| **Y128F** | $1.74±0.19·10^{-1}$ |
| **Y128A** | $3.13±0.10·10^{-1}$ |
| **Y128E** | $1.03±0.01·10^{-1}$ |

**Table S3. Mid-point reduction potentials for NQO1 variants.** Potentials determined by using the Xanthine/XO method in 20 mM HEPES-KOH pH 7.4 at 25 °C. Examples for primary data and fittings are shown in Figure S15.

| NQO1 variant | $E_{ox/hq}$ (mV) |
|---|---|
| **WT** | -124.6 ± 3.1 |
| **Y126A** | -135.7 ± 0.7 |
| **Y128A** | -132.0 ± 0.8 |
| **Y126E** | -149.3 ± 0.3 |
| **Y128E** | -128.1 ± 0.1 |
| **Y126F** | -133.4 ± 1.3 |
| **Y128F** | -137.5 ± 2.9 |

**Supplementary References**

1 Medina-Carmona E, Fuchs JE, Gavira JA, Mesa-Torres N, Neira JL, Salido E, Palomino-Morales R, Burgos M, Timson DJ & Pey AL (2017) Enhanced vulnerability of human proteins towards disease-associated inactivation through divergent evolution. *Hum Mol Genet* 26, 3531-3544.

2 Vankova P, Salido E, Timson DJ, Man P & Pey AL (2019) A Dynamic Core in Human NQO1 Controls the Functional and Stability Effects of Ligand Binding and Their Communication across the Enzyme Dimer. *Biomolecules* 9, 728.
